# Effects of atmospheric CO_2_ levels on the susceptibility of maize to diverse pathogens

**DOI:** 10.64898/2025.12.31.697224

**Authors:** Ekkachai Khwanbua, Yunhui Qi, John Ssengo, Peng Liu, Michelle A. Graham, Steven A. Whitham

**Affiliations:** Department of Plant Pathology, Entomology and Microbiology, Iowa State University, Ames, IA 50011, USA; Department of Statistics, Iowa State University, Ames, IA 50011, USA; Department of Data Science, Dana-Farber Cancer Institute, Boston, MA 02215, USA; Interdepartmental Genetics and Genomics, Iowa State University, Ames, IA 50011, USA; United States Department of Agriculture (USDA), Agricultural Research Service (ARS), Corn Insects and Crop Genetics Research Unit and Department of Agronomy, Iowa State University, Ames, IA 50011, USA

**Keywords:** *Zea mays*, maize, carbon dioxide, plant immunity, plant defense, PAMP-triggered immunity, disease susceptibility, basal defense

## Abstract

Rising atmospheric CO_2_ has profound implications for crop productivity and food security. Based on studies in C3 plants, elevated CO_2_ (eCO_2_) can shape plant-pathogen interactions, although the outcomes are often variable. The question of how eCO_2_ influences immunity and disease development in C4 plants, such as the globally important cereal crop maize (*Zea mays* L.), has not been systematically examined. We challenged maize plants grown under ambient CO_2_ (aCO_2_, 420 ppm) and eCO_2_ (550 ppm) with bacterial, viral, fungal, and oomycete pathogens. Plants grown in eCO_2_ were more susceptible to sugarcane mosaic virus, suggesting compromised antiviral defenses, less susceptible to *Clavibacter nebraskensis*, *Exserohilum turcicum*, and *Colletotrichum graminicola*, and susceptibility to *Puccinia sorghi* and *Pythium sylvaticum* was unchanged. Reduced susceptibility to *C. nebraskensis* was associated with enhanced basal immune responses. These results establish a foundation for dissecting eCO_2_-responsive defense mechanisms, and they highlight a critical need to understand how eCO_2_ will impact plant responses to microbes, pests, and abiotic stresses under future conditions.

## Main

The concentration of carbon dioxide (CO_2_) in the atmosphere has risen steadily in proportion to human activities involving fossil fuel combustion and land use changes since the Industrial Revolution^1^. Unlike other abiotic factors that vary across space and time, CO_2_ is well-mixed in the atmosphere and continues to rise globally^2^. At current emission rates, atmospheric CO_2_ levels are expected to reach 550 parts per million (ppm) by 2050-2100^3^. In C3 plants, elevated CO_2_ (eCO_2_) generally enhances photosynthesis^4^ and water use efficiency^5^, resulting in greater biomass and yield^6^. However, eCO_2_ also alters leaf chemical composition, such as nonstructural carbohydrate^7^, nitrogen and protein^8^, while also triggering changes in secondary metabolism^9^, defense responses^10^, chemical signaling^11^, and root exudation^12^. In C4 plants, photosynthesis is not stimulated in eCO_2_, because CO_2_ is concentrated in bundle sheath cells where photosynthesis occurs^6^, and they tend to maintain consistent foliar nitrogen and secondary metabolite levels. Similar to C3 plants, water use efficiency is increased in C4 plants in eCO_2_ due to decreased stomatal apertures^13^.

Plant disease results from interactions among a susceptible host, a virulent pathogen, and the environment^14^. Therefore, shifts in environmental variables, including rising CO_2_ levels, are likely to affect plant-pathogen interactions^15^. In studies conducted almost exclusively in C3 plants, eCO_2_ is associated with changes in plant defense responses, including plant architecture^16^, stomatal density and aperture^17^, redox regulation^18^, mitogen activated protein kinase (MAPK) expression^19^, secondary metabolites^9^, and phytohormone production^20^. In general, plants grown under eCO_2_ exhibit increased constitutive and induced salicylic acid (SA) levels while suppressing jasmonate levels^10^, suggesting enhanced resistance to biotrophic pathogens and greater susceptibility to necrotrophic pathogens. However, the degree and directionality of these responses often vary, which may reflect species- or genotype-specific responses to eCO_2_^10^. The variations among studies emphasize the need for direct investigations into each pathosystem involving important crop species, rather than depending on generalizations from model systems.

Although C4 plants comprise a small proportion of the world’s plant species, they contribute 18-21% of global primary productivity^21^. Maize (*Zea mays* L.) is the most widely grown annual cereal C4 crop and is used for food, feed, and biofuel^22^, accounting for approximately 1.23 billion metric tons of global production in 2023 (http://www.worldagriculturalproduction.com/). While extensive research has assessed the effects of eCO_2_ on C3 plants and their pests, the impacts of eCO_2_ on C4 plant-pathogen interactions are virtually unexplored. In particular, two studies have investigated the effects of eCO_2_ on maize susceptibility to *Fusarium verticillioides* and its mycotoxin^23,24^. Given that maize production is affected by many economically important pathogens^25^, a comprehensive study of how eCO_2_ affects maize susceptibility to diverse pathogens is needed. Moreover, the effects of eCO_2_ on defense responses have yet to be examined at the molecular level in this and other C4 plant species.

In this study, we compared the impacts of current aCO_2_ (420 ppm) and eCO_2_ (550 ppm) on maize immune responses and disease susceptibility using phytopathogens chosen for their diverse infection strategies and relevance to important maize diseases. We assessed basal immunity using the maize-*Clavibacter nebraskensis* pathosystem and performed transcriptomic analyses on this pathosystem and on sugarcane mosaic virus and *Puccinia sorghi* to gain an in-depth understanding of how eCO_2_ affects these interactions. We also conducted infection experiments using *Exserohilum turcicum*, *Colletotrichum graminicola*, and *Pythium sylvaticum.* Our work demonstrates that eCO_2_ has diverse effects on maize interactions with these microbial pathogens and establishes a basis for future investigations into the molecular mechanisms that govern defense responses in eCO_2_ environments, while pinpointing pathogens of potential concern in predicted future atmospheric scenarios.

## Results

### eCO_2_ induces physiological changes in maize

Prior to investigating how eCO_2_ affects maize susceptibility to plant pathogens, we first evaluated whether our growth conditions produced physiological responses commonly associated with eCO_2_. Plants grown in eCO_2_ exhibited lower stomatal conductance to water vapor (g_sw_) and transpiration (E) compared to plants grown in aCO_2_ (Fig. 1a,b), consistent with decreased stomatal aperture (Fig. 1c) and no change in stomatal density on the abaxial leaf surface (Fig. 1d). Photosystem II (ΦPSII) efficiency did not differ between plants grown at the two different CO_2_ levels (Fig. 1e), indicating no increase in photosynthetic rate. At 42 days after planting (dap), maize plants grown in eCO_2_ had greater fresh weight (Fig. 1f), dry weight (Fig. 1g), and height (Fig. 1h) than plants grown in aCO_2_ (Fig. 1i). Together, these experiments suggest that the future predicted elevated atmospheric [CO_2_] of 550 ppm alters physiological responses in maize that align with previous studies^26,27^.

**Fig. 1.**
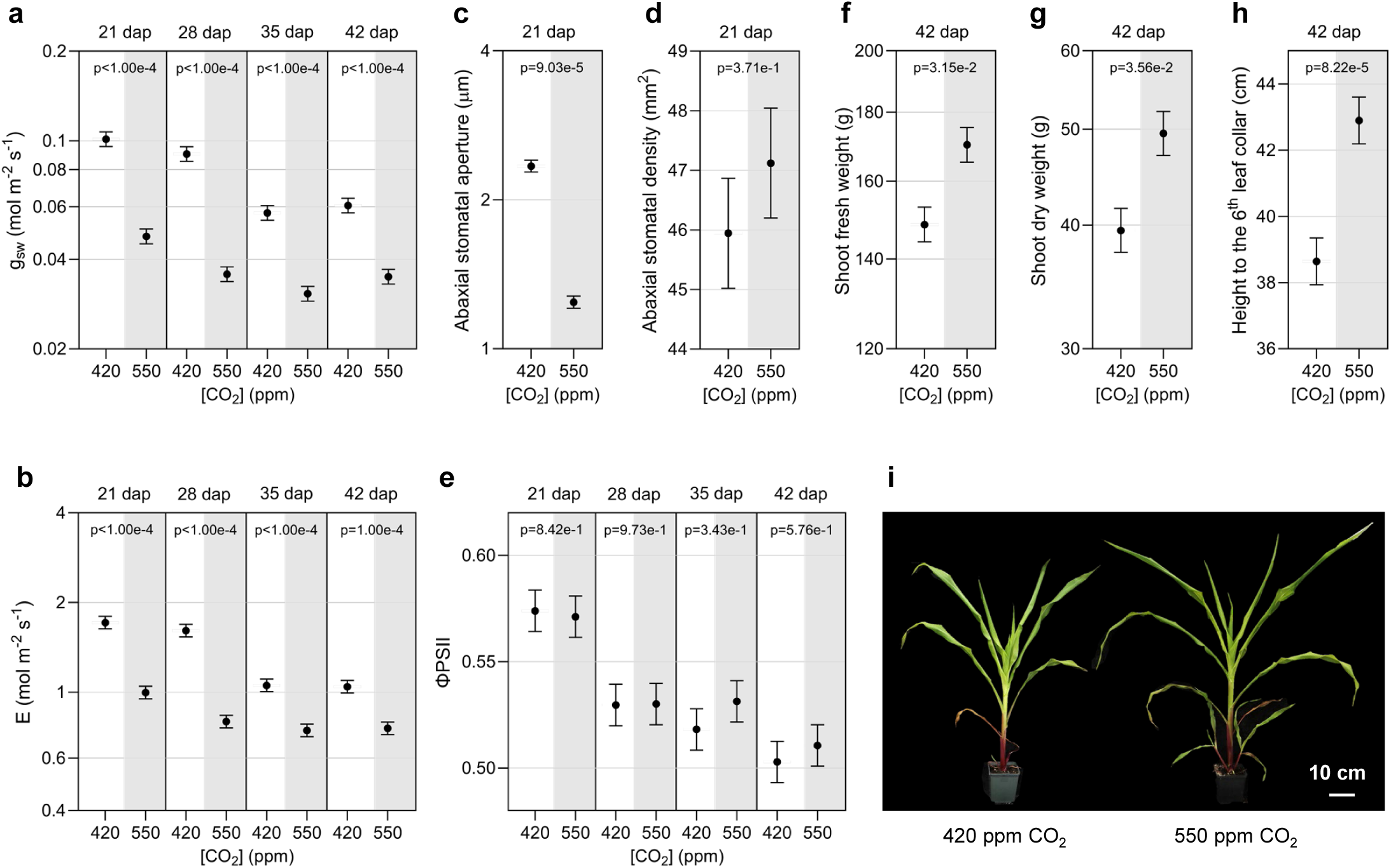
Impact of elevated CO_2_ on maize physiological traits. **a-b**, Stomatal conductance (g_sw_) and transpiration (E) were assessed at the indicated days after planting (dap) using the LI-600 system. **c-d**, Abaxial stomatal aperture and stomatal density were quantified at 21 dap using light microscopy, with three randomly selected fields of view per leaf sample. **e**, photosystem II (ΦPSII) efficiency was assessed at the indicated dap using the LI-600 system. **f-h**, Shoot fresh weight, shoot dry weight, and plant height measured to the sixth leaf collar were recorded at 42 dap. **i**, Representative photos of maize plants were taken at 42 dap. Measurements and samples were taken on the fifth fully expanded leaf at 21 and 28 dap and on the sixth fully expanded leaf at 35 and 42 dap, using ten plants per time point. The three experimental replicates were conducted simultaneously in independent CO_2_ controlled chambers using a completely randomized design. Data were graphed as the mean across the three replicates with standard error (SE) bars. For variables with unequal variance (g_sw_, E, abaxial stomatal aperture, shoot fresh weight, and shoot dry weight), linear mixed effect model (LMM) analysis was applied to the log-transformed data, and means with SE are presented on the log10 scale. *P*-values for g_sw_, E, and ΦPSII were derived from pairwise contrasts using *t*-tests, whereas those for abaxial stomatal density, abaxial stomatal aperture, shoot fresh weight, and plant height were obtained from *F*-tests of the main CO_2_ effect from the LMM analysis. The letter e denotes an exponent to the power of 10 and ppm denotes parts per million.

### Maize is less susceptible to *C. nebraskensis* in eCO_2_

To determine if eCO_2_ affects maize susceptibility to *C. nebraskensis,* the bacterium causing Goss’s wilt and leaf blight^28^, we inoculated the third leaf of two-week-old plants using a leaf-tip clipping method. *C. nebraskensis*-infected maize plants grown in aCO_2_ displayed earlier blight symptoms and greater lesion lengths at 4 and 8 dpi relative to plants grown in eCO_2_ (Fig. 2a-c). Consistent with disease development, less bacterial multiplication was observed in maize leaf tissue at 5 cm and 10 cm from the inoculation sites at 4 and 8 dpi in eCO_2_ (Fig. 2d,e). To assess if eCO_2_ directly affects *C. nebraskensis* growth rate, bacterial growth was assessed in liquid cultures in the CO_2_-controlled growth chambers. Bacterial growth was nearly identical between the two [CO_2_] conditions (Supplementary Fig. 1), suggesting reduced bacterial growth did not contribute to the diminished growth *in planta* in eCO_2_.

**Fig. 2.**
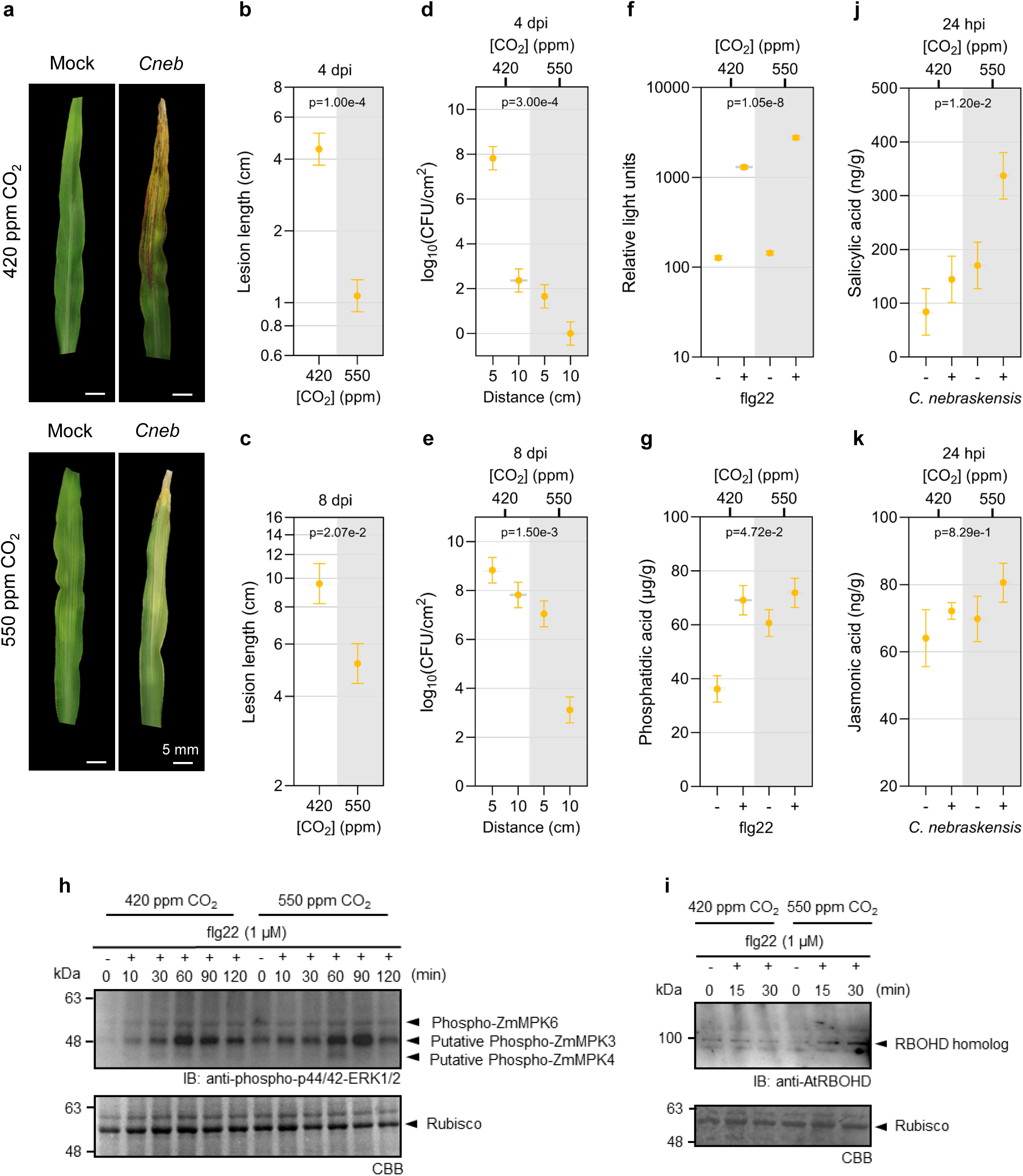
Elevated CO_2_ reduces maize susceptibility to *Clavibacter nebraskensis* (*Cneb*) and enhances bacterially-induced immune signaling. **a,** Representative images of the third maize leaf inoculated with *C. nebraskensis* or mock treated with phosphate-buffered saline were photographed at 8 days post inoculation (dpi). **b-e**, Lesion lengths and bacterial populations on the third leaf, quantified as colony-forming units (CFUs) at 5 cm and 10 cm from the inoculation site, were measured at 4 and 8 dpi. Six plants were analyzed per CO_2_ treatment at each time point. **f**, Reactive oxygen species (ROS) production in response to flg22 was quantified using a chemiluminescence assay. Relative light units were determined using leaf discs from 12 plants per CO_2_ treatment. **g**, Phosphatidic acid (PA) accumulation was quantified using liquid chromatography-tandem mass spectrometry (LC-MS/MS) following a 5-minute incubation with the bacterial peptide flg22 or a mock treatment. For each CO_2_ condition, leaf discs were collected separately from 6 flg22-treated and 6 mock-treated plants, pooled within each treatment group to form one biological sample, and three biological samples were analyzed per treatment. **h**, MAPK activation following flg22 treatment was assessed by immunoblot using protein extracted from 6 plants per treatment for each CO_2_ condition at 0 (untreated), 10, 30, 60, 90, and 120 min after treatment. **i**, Accumulation of RBOHD was analyzed by immunoblot using leaf tissues from mock- and flg22-treated plants, with 6 plants per treatment at each CO_2_ level. Samples were collected at 0 (untreated), 15, and 30 min after treatment. **j-k**, Salicylic acid (SA) and jasmonic acid (JA) accumulation at 24 hpi in response to *C. nebraskensis* infection was quantified using targeted LC-MS/MS analysis with six plants per treatment at each CO_2_ concentration.The three experimental replicates were carried out simultaneously in independent CO_2_-controlled chambers using a replicated completely randomized design. Data were graphed as the mean across the three replicates with SE bars. Due to unequal variance on the original scale, linear mixed effect model (LMM) was applied to the log-transformed lesion length and ROS data, and means with SE are presented on the log10 scale. LMM with different residual variances across different CO_2_-by-pathogen combinations was applied to JA data to account for heteroskedasticity. *P*-values for lesion length and CFUs were obtained from pairwise contrasts using *t*-tests. *P*-values for ROS and PA were obtained from *F*-tests of the interaction effect between CO_2_ and pathogen treatment. *P*-value for JA was obtained from Wald *Chi*-square tests of the interaction effect between CO_2_ and pathogen treatment, and the *p*-value for SA was obtained from an *F*-test of the main CO_2_ effect. The letter e denotes an exponent to the power of 10 and ppm denotes parts per million.

### eCO_2_ affects basal immune responses

To assess whether eCO_2_ alters basal maize defenses during *C. nebraskensis* infection, we examined early hallmarks of pattern-triggered immunity (PTI). Upon recognition of microbe associated molecular patterns (MAMPs) by plasma membrane localized pattern recognition receptors (PRRs), plants trigger reactive oxygen species (ROS) production and phosphorylation cascades^29^, and activate phospholipid modifying enzymes such as phospholipase D (PLD) and diacylglycerol kinases (DGKs) to generate phosphatidic acid (PA). PA helps stabilize respiratory burst oxidase homolog D (RBOHD) and sustain the ROS burst^30^. Maize plants grown in eCO_2_ exhibited stronger ROS production (Fig 2f) in response to flg22 and greater constitutive PA levels (p-value = 0.0047) (Fig 2g) compared to plants grown under aCO_2_. We also observed greater MAPK activation (Fig 2h) and a modest increase in maize RBOHD protein accumulation (Fig. 2i, Supplementary Fig. 2). In addition, plants grown in eCO_2_ accumulated higher levels of salicylic acid (SA) at 24 hpi with *C. nebraskensis* relative to plants in aCO_2_ (Fig. 2j), whereas jasmonic acid (JA) levels did not differ between the two CO_2_ concentrations (Fig. 2k). Together, these data suggest that plants grown at eCO_2_ mounted a more robust immune response to the flg22 peptide and bacterial challenge, consistent with enhanced resistance to *C. nebraskensis*.

### Maize susceptibility to SCMV is enhanced in eCO_2_

To investigate whether eCO_2_ impacts maize susceptibility to SCMV, we mechanically inoculated the seventh leaf of four-week-old plants^31^. At 21 dpi, the leaves of SCMV-infected plants grown in eCO_2_ exhibited more pronounced mosaic symptoms compared to those in aCO_2_, with more widespread areas of light and dark green mosaic symptoms (Fig. 3a). Despite no difference in plant height between SCMV-infected plants under eCO_2_ and aCO_2_ compared to mock plants (Supplementary Fig. 3a), SCMV caused a larger reduction in shoot dry weight in eCO_2_ than in aCO_2_ relative to mock-treated plants, but fresh weight was not significantly affected (Fig. 3b, Supplementary Fig. 3b). The accumulation of SCMV in the newest fully expanded leaves of infected plants was significantly higher in eCO_2_ at both 14 and 21 dpi (Fig. 3c, Supplementary Fig. 3c), consistent with the greater area under disease progress curve (AUDPC) for SCMV-infected plants grown in eCO_2_ (Fig. 3d). At 14 and 21 dpi, SCMV-infected plants grown in eCO_2_ showed a greater reduction in stomatal conductance and transpiration than those in aCO_2_, relative to mock-inoculated plants, although this difference was less pronounced at 21 dpi (Fig. 3e,f, Supplementary Fig. 3d,e). To further test if stomatal aperture or density were affected by SCMV, we imaged epidermal leaf impressions collected at 14 dpi using a scanning electron microscope. Maize plants grown at eCO_2_ exhibited a smaller overall stomatal aperture when considering both mock- and SCMV-inoculated plants (Fig. 3g), and stomatal density was not significantly affected (Fig. 3h). As observed in the mock-inoculated plants, photosynthetic efficiency in SCMV-infected plants did not differ between aCO_2_ and eCO_2_ at either 14 or 21 dpi (Fig. 3i, Supplementary Fig. 3f).

**Fig. 3.**
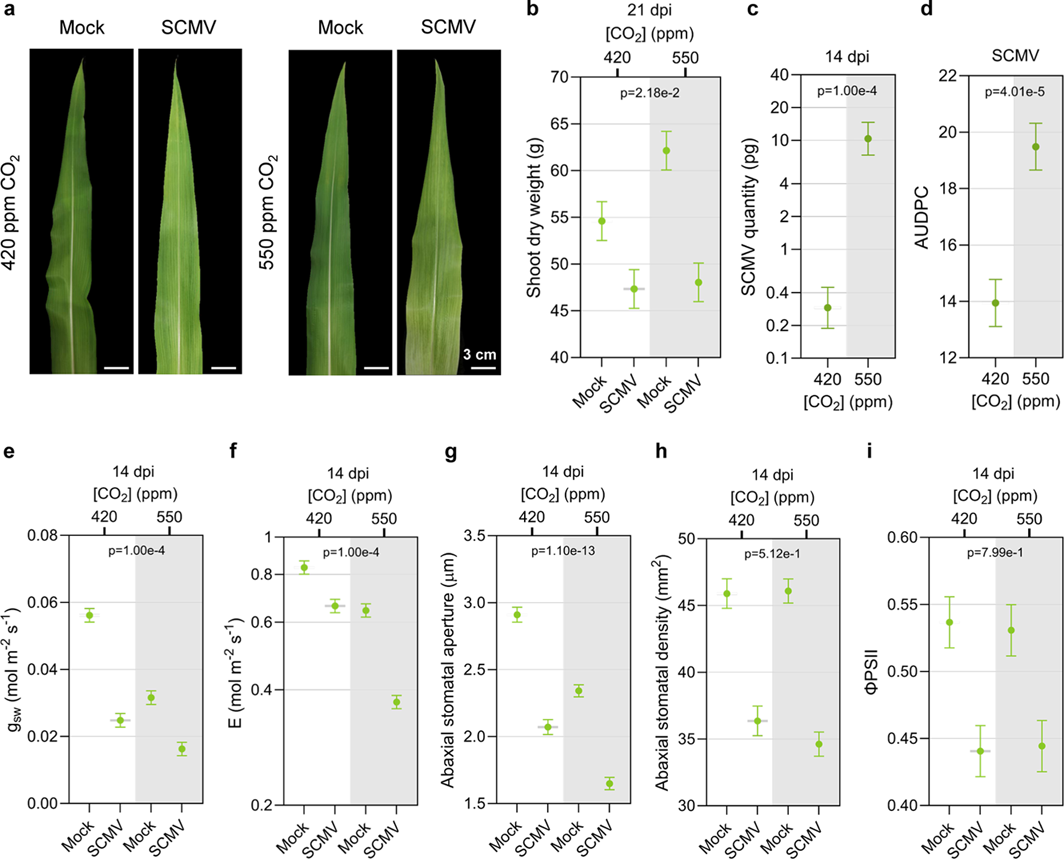
Maize plants grown under elevated CO_2_ are more susceptible to sugarcane mosaic virus (SCMV). **a**, Representative photos of the twelfth maize leaf in response to ambient CO_2_ (aCO_2_) or elevated CO_2_ (eCO_2_) during SCMV infection, compared with mock-treated plants, photographed at 21 days post inoculation (dpi). **b**, Shoot dry weight of maize plants was measured at 21 dpi following SCMV or mock treatment. **c**, SCMV accumulation was quantified by reverse transcriptase-quantitative PCR (RT-qPCR) using SCMV-specific primers. Absolute quantification was performed using a standard curve generated from serial dilutions of the SCMV infectious clone. **d**, SCMV disease severity quantified as the area under the disease progress curve (AUDPC) over a 49-day time course. **e-f**, Stomatal conductance (g_sw_) and transpiration rate (E) were measured on the abaxial side of the eleventh leaf of mock- and SCMV-treated maize plants at 14 dpi using the LI-600 system. **g-h**, Stomatal aperture and density were assessed on the abaxial leaf surface at 14 dpi using a scanning electron microscope. For both measurements, six plants were used per treatment and CO_2_ condition. Five fields of view were randomly selected per plant for stomatal aperture, and two fields of view per plant were used for stomatal density. **i,** Photosystem II efficiency (ΦPSII) was measured on the adaxial side of the eleventh leaf of mock- and SCMV-treated maize plants at 14 dpi using the LI-600 system. The three experimental replicates were conducted simultaneously in independent CO_2_-controlled chambers under a replicated completely randomized design. Data are shown as means across the three replicates with SE bars. Due to unequal variance on the original scale, linear mixed effect model (LMM) was applied to the log-transformed SCMV quantity and E data, and means with SE are presented on the log10 scale. *P*-values for AUDPC, abaxial stomatal aperture, and abaxial stomatal density were obtained from *F*-tests of the main CO_2_ effect, while that for shoot dry weight was obtained from *F*-tests of interaction effects between CO_2_ and pathogen treatment. *P*-value for SCMV quantity was obtained from the pairwise *t*-test comparing the two CO_2_ levels at 14 dpi, whereas *p*-values for g_sw_, E, and ΦPSII were from the *t*-tests of the interaction effect between CO_2_ and pathogen treatment at 14 dpi from the LMM analyses. The letter e denotes an exponent to the power of 10, and ppm denotes parts per million.

### eCO_2_ does not alter maize susceptibility to common rust

To determine whether eCO_2_ affects maize susceptibility to common rust caused by the obligate biotrophic fungus *P. sorghi*^32^, we first considered that *P. sorghi* requires stomatal entry to initiate infection^33^. Given that stomatal traits can influence pathogen entry, we measured stomatal conductance and aperture at 10 dap, just prior to inoculation. As expected, the non-inoculated maize plants grown under eCO_2_ exhibited significantly lower stomatal conductance (Fig. 4a) and narrower stomatal aperture (Fig. 4b,c). We then spray-inoculated a spore suspension onto the third and fourth leaves of ten-day-old maize plants. *P. sorghi* successfully established infection and formed pustules on maize leaves under both [CO_2_] (Fig. 4d). While the mean pustule size of *P. sorghi*-infected plants grown in eCO_2_ was greater than those grown in aCO_2_ (Fig. 4e), no difference was observed in pustule coverage and number between *P. sorghi*-infected plants grown under eCO_2_ and aCO_2_ at 8 dpi (Fig. 4f,g), which was consistent with similar levels of rust accumulation at 18 hpi, 3 dpi, and 5 dpi (Fig. 4h).

**Fig. 4.**
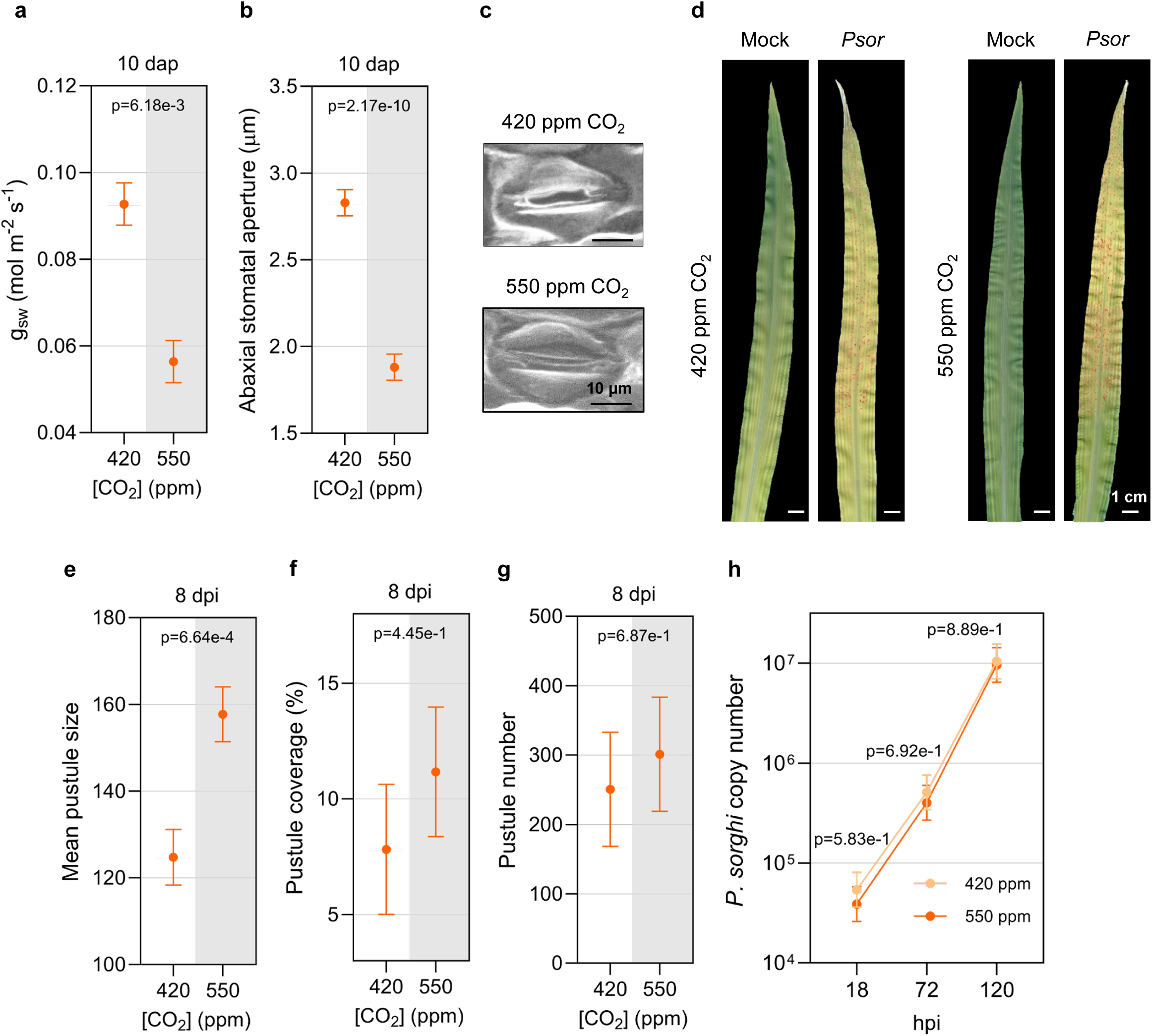
Elevated CO_2_ does not alter maize susceptibility to *Puccinia sorghi* (*Psor*). **a-b**, Stomatal conductance (g_sw_) and aperture were measured on the abaxial surface of the third leaf prior to inoculation, using six plants per CO_2_ treatment. Aperture measurements were based on five randomly selected fields of view per leaf sample, imaged using scanning electron microscopy. **c**, Close-up images of maize stomata on the abaxial surface of the third leaf under aCO_2_ or eCO_2_, taken at 10 days after planting (dap). **d**, Representative images of the third maize leaf under ambient CO_2_ (aCO_2_) or elevated CO_2_ (eCO_2_), either mock-treated or infected with *P. sorghi*, photographed at 8 days post inoculation (dpi). **e-g**, Mean pustule size, pustule coverage, and pustule number were quantified from eight *P. sorghi*-infected plants per CO_2_ treatment, collected at 8 dpi, using a U-Net neural network trained for the *P. sorghi*-maize pathosystem. **h**, For fungal biomass quantification, leaf tissue from two *P. sorghi*-infected maize plants was pooled to form one sample. At each time point (18, 72, 120 hours post inoculation (hpi)), eight plants per treatment under each CO_2_ level were sampled, resulting in three independent biological replicates per treatment. Fungal DNA was quantified by quantitative polymerase chain reaction (qPCR) using *P. sorghi*-specific primers, and absolute quantification was performed using a standard curve generated from serial dilutions of known fungal DNA concentrations. The three experimental replicates were conducted simultaneously in independent CO_2_-controlled chambers using a replicated complete randomized design. Data were graphed as the mean across the three replicates with SE bars. Due to unequal variance on the original scale, linear mixed effect model was applied to the log-transformed DNA copy number data, and means with SE are presented on the log10 scale. *P*-values for g_sw_, abaxial stomatal aperture, pustule number, percent pustule coverage, and mean pustule size were obtained from *F*-tests of the main CO_2_ effect. *P*-values for DNA copy number were derived from pairwise contrasts (*t*-tests) of the between the two CO_2_ levels. The letter e denotes an exponent to the power of 10, and ppm denotes parts per million.

### Maize is less susceptible to *E. turcicum* and *C. graminicola* in eCO_2_

To investigate the effects of eCO_2_ on maize susceptibility to the hemibiotrophic fungus *E. turcicum* (northern corn leaf blight (NCLB))^34^, we inoculated infested millet into the whorls of three-week-old maize plants. The infected plants grown in eCO_2_ exhibited moderate NCLB symptoms, whereas those grown in aCO_2_ developed more severe symptoms (Fig. 5a). The reduced NCLB symptoms in eCO_2_ corresponded to a slower progression of foliar symptoms (Fig. 5b) and lower *E. turcicum* accumulation (Fig. 5c) in infected plants grown under eCO_2_ at 28 dpi. We assessed the expression of maize homologs of two genes associated with defense responses to pathogen infections, *ZmPR1* (SA marker gene) and *ZmLOX9* (JA marker gene). The expression of *ZmPR1* in infected plants was higher under eCO_2_ compared to aCO_2_, relative to mock-inoculated plants (Fig. 5d), while no difference was observed in *ZmLOX9* expression (Supplementary Fig. 4a).

**Fig. 5.**
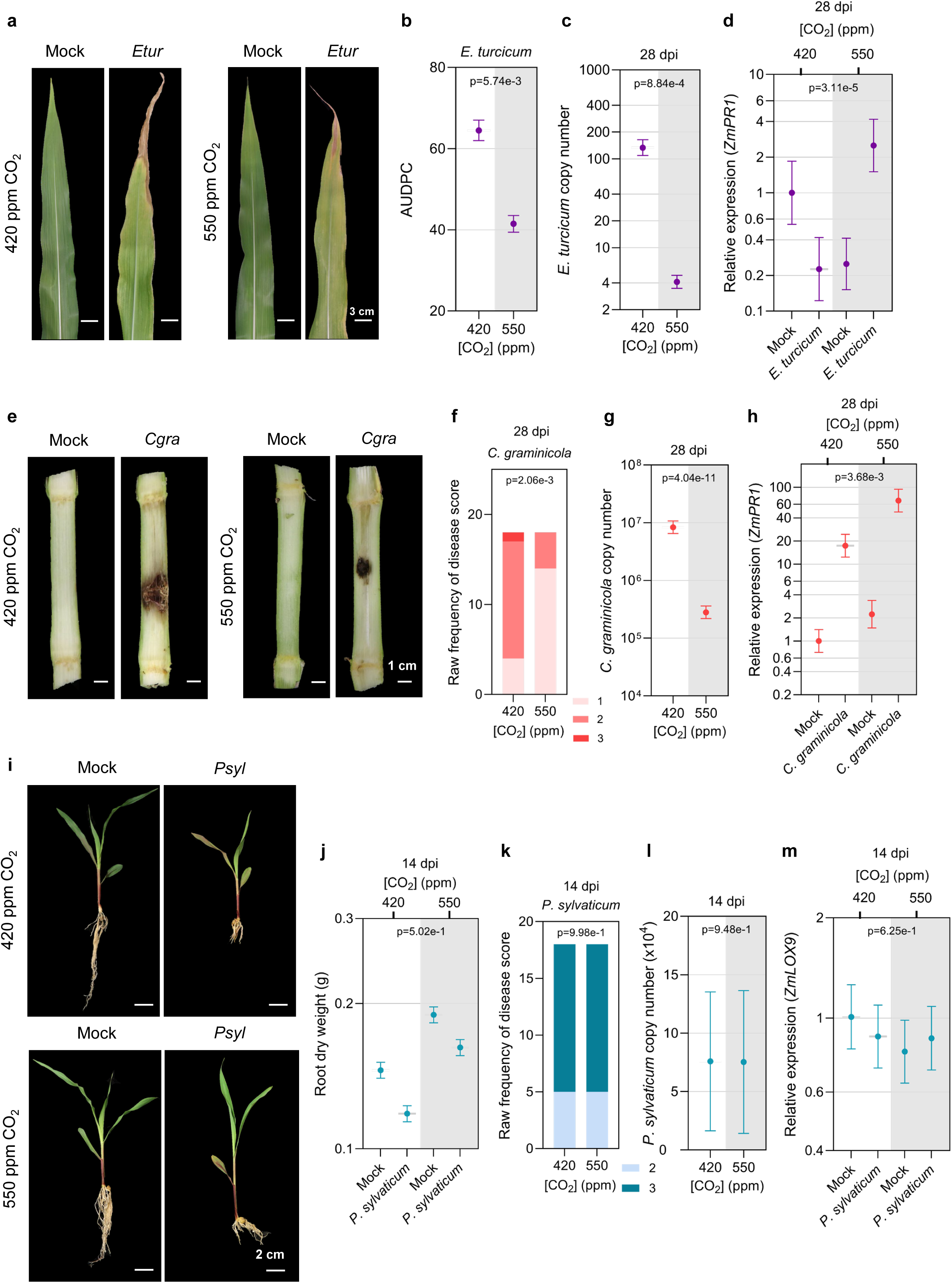
Differential effects of elevated CO_2_ on maize susceptibility to filamentous pathogens. **a**, Representative photos of the seventh maize leaf under ambient CO_2_ (aCO_2_) or elevated CO_2_ (eCO_2_) that were mock-treated or inoculated with *Exserohilum turcicum* (*Etur*) photographed at 28 days post inoculation (dpi). **b**, Northern leaf blight severity quantified as the area under the disease progress curve (AUDPC) over a 49-day time course. **c**, Quantification of *E. turcicum* was performed by quantitative PCR (qPCR) using pathogen-specific primers, and absolute values were determined from a standard curve generated from known concentrations of fungal DNA. **d**, Relative expression of *ZmPR1* (SA marker gene) in maize was assessed by reverse transcriptase quantitative polymerase chain reaction (RT-qPCR) analysis relative to the *Zmβ-TUB*. **e**, Representative images of maize stalks under aCO_2_ or eCO_2_ that were mock-treated or infected with *Colletotrichum graminicola* (*Cgra*), photographed at 28 dpi. **f**, *C. graminicola* raw frequency of disease score based on a 0-5 scale where 0 indicates no visible symptoms and 5 indicates complete stalk necrosis with tissue collapse. Only scores 1–3 were observed in this experiment. Intermediate scores reflect increasing levels of necrosis and tissue degradation. **g**, *C. graminicola* quantity was quantified by qPCR with pathogen-specific primers, using a standard curve generated from known concentrations of fungal DNA for absolute quantification. **h**, Relative expression of *ZmPR1* (SA marker gene) in maize was assessed by RT-qPCR analysis relative to the *Zmβ-TUB*. **i**, Representative images of maize plants grown in aCO_2_ or eCO_2_ at 14 days after planting (dap) that were mock-treated or germinated in soil infested with *Pythium sylvaticum* (*Psyl*). **j**, Root dry weight of maize plants measured at 14 dap following mock or *P. sylvaticum* treatment. **k**, *P. sylvaticum* disease severity at 14 dap, shown as raw frequency of severity scores. Disease was assessed using a 0-4 scale based on root condition, where 0 indicates long, healthy roots with no discoloration; 1 indicates slightly stunted and moderately discolored roots; 2 indicates severe stunting and/or discoloration; 3 indicates short, rotted roots; and 4 indicates rotted seed with no germination. Only scores 2 and 3 were observed under the conditions tested. **l**, *P. sylvaticum* quantity assessed by qPCR using species-specific primers, with absolute values calculated from a standard curve prepared from known concentrations of oomycete DNA. **m**, Relative expression of *ZmLOX9*, a jasmonic acid (JA) marker gene, was assessed by RT-qPCR relative to *Zmβ-TUB*. Three experiments were conducted simultaneously in independent CO_2_ control chambers, using 6 plants per treatment for each CO_2_ concentration according to a replicated complete randomized design. Data were graphed as the mean across the three replicates with SE bars. For *E. turcicum*, due to unequal variance on the original scale, linear mixed effect model (LMM) was applied to the log-transformed DNA copy number and *ZmPR1* expression, and means with SE are presented on the log10 scale. *P*-values for AUDPC and DNA copy number were obtained from *F*-tests of the main CO_2_ effect, and those for *ZmPR1* expression from *F*-tests of the interaction effect from the LMM analyses. For *C. graminicola*, cumulative link mixed model (CLMM) was applied to disease scores and *p*-value was obtained from the likelihood ratio tests, while DNA copy number and *ZmPR1* expression were tested with *F*-tests of the main CO_2_ effect on log-transformed data from the LMM analyses. For *P. sylvaticum*, the *P*-value for DNA copy number was obtained from the *F*-test of the main CO_2_ effect in LMM analysis. The *P*-value for disease scores was obtained from the likelihood ratio test in CLMM analysis, and the *p*-value for *ZmLOX9* expression was from the *F*-test of the interaction effect in the LMM analysis of the log-transformed data. The letter e denotes an exponent to the power of 10, and ppm denotes parts per million.

To assess whether eCO_2_ affects maize susceptibility to the hemibiotrophic fungus *C. graminicola* causing stalk rot^35^, we inoculated the stalks of six-week-old plants using sterilized toothpicks soaked in a spore suspension. At 28 dpi, *C. graminicola*-infected plants grown at eCO_2_ exhibited less severe disease than plants grown at aCO_2_ (Fig. 5e). The decreased disease severity in eCO_2_ was consistent with a lower disease index (Fig. 5f) and reduced *C. graminicola* accumulation (Fig. 5g) in the stalk tissue compared to plants grown under aCO_2_. To investigate whether eCO_2_ affected the expression of SA-and JA-related defense genes, *ZmPR1* and *ZmLOX9* mRNA transcript accumulation was quantified. *ZmPR1* transcript levels were higher under eCO_2_ when considering mock- and *C. graminicola*-inoculated plants (Fig 5h), while *ZmLOX9* expression showed no significant difference under the two CO_2_ conditions (Supplementary Fig. 5a).

To determine whether *E. turcicum* or *C. graminicola* growth *in vitro* is affected by [CO_2_], we measured the diameter of fungal colonies on plates maintained under eCO_2_ or aCO_2_ every other day for 15 days. The growth of each fungus was similar under the two [CO_2_] conditions, suggesting that eCO_2_ may alter maize susceptibility by modulating host defense responses rather than by directly affecting pathogen growth *in planta* (Supplementary Fig. 4b, 5b).

### Maize susceptibility to *P. sylvaticum* is unchanged in eCO_2_

To determine whether eCO_2_ affects maize susceptibility to the soil-borne necrotroph *P. sylvaticum* (seedling blight and root rot)^36^, we germinated kernels in sand-soil mixed with infested millet. While *P. sylvaticum* caused noticeable root rot disease under both CO_2_ conditions (Fig. 5i), shoot and root fresh weights (Supplementary Fig. 6a,b) and shoot and root dry weights (Supplementary Fig. 6c, Fig. 5j) remained similar between *P. sylvaticum*-infected and mock-treated plants under eCO_2_ relative to aCO_2_ at 14 dap. However, increases in root fresh weight and root dry weight of mock-inoculated plants grown at eCO_2_ compared to those at aCO_2_ were observed at 14 dap (p-values = 0.0094 and 0.0012, respectively). The disease index indicated similar disease severity between *P. sylvaticum*-infected plants grown under both CO_2_ treatments (Fig. 5k), consistent with the *P. sylvaticum* titer measured in the roots at 14 dap (Fig. 5l). No difference was observed in *ZmPR1* (Supplementary Fig. 6d) and *ZmLOX9* (Fig. 5m) expression in response to *P. sylvaticum* under both CO_2_ conditions. In addition, seed rot severity of *P. sylvaticum*-infected seeds was similar between the two CO_2_ treatments (p-value = 0.1519) (Supplementary Fig. 6e), despite the faster growth rate of *P. sylvaticum* at eCO_2_^17^.

### Maize transcriptome responses to *C. nebraskensis*, SCMV, and *P. sorghi* are differentially modulated by eCO_2_

To investigate how *C. nebraskensis*, SCMV, and *P. sorghi* modulate host gene expression in response to different [CO_2_], we performed QuantSeq 3′ mRNA-Seq on leaf tissues treated with mock (buffer only) or pathogen inoculation. We identified DEGs responding to pathogen infection at different [CO_2_] (Path aCO_2_ vs Mock aCO_2_ and Path eCO_2_ vs Mock eCO_2_, FDR < 0.05, Supplementary Tables 6, 8, 10, 12 and 14) and DEGs responding to [CO_2_] in pathogen- or mock-inoculated tissues (Path eCO_2_ vs Path aCO_2_ and Mock eCO_2_ vs Mock aCO_2_, FDR < 0.05, Supplementary Tables 7, 9, 11, 13 and 15). For each DEG list, we used GO term enrichment to confirm expected responses to pathogen inoculation and eCO_2_ (Supplementary Table 16). In addition, clustering and heat map visualization confirmed defense and eCO_2_ responses (Supplementary Fig. 7-11).

To identify DEGs that were pathogen and eCO_2_ responsive, we selected significant DEGs expressed in response to pathogen inoculation in aCO_2_ and/or eCO_2_ and then retained only those DEGs that were also significant for their CO_2_ response in either mock and/or pathogen-treated samples. This filtering identified 2,154 DEGs responsive to *C. nebraskensis* and [CO_2_] at 4 dpi, 147 DEGs responsive to SCMV and [CO_2_] at 14 dpi, and 8,106, 1,865 and 4,711 DEGs responsive to *P. sorghi* and [CO_2_] at 18, 72 and 120 hpi, respectively. Combining across pathogen treatments resulted in 11,640 unique pathogen and eCO_2_ responsive DEGs. Using this gene set, we generated two heat maps of Z score normalized log_2_ fold change values. The first heat map (Fig. 6a, Supplementary Table 17) reveals how pathogen-induced expression patterns differ between aCO_2_ and eCO_2_. This heat map demonstrates the diversity of responses to pathogen infection. Following *C. nebraskensis* infection, the magnitude of pathogen responsiveness in all clusters is greater in aCO_2,_ suggesting that this pattern reflects the greater bacterial accumulation in aCO_2_. For SCMV, pathogen-responsiveness is similar under both CO_2_ conditions, suggesting eCO_2_ has little impact on responses to SCMV. However, SCMV-infected plants grown under eCO_2_ showed strong downregulation of two *AGO1* and two *AGO10* homologs (Supplementary Table 8), which is consistent with the higher SCMV accumulation in eCO_2_. As AGOs are essential for executing RNA silencing, their reduced expression suggests a possible mechanism for weakened antiviral defenses. For *P. sorghi*, we see mixed expression patterns across time points and in response to eCO_2_. The second heat map (Fig. 6b, Supplementary Table 18) displays how pathogen infection modifies the plant transcriptional response to eCO_2_. Infection with *C. nebraskensis* increases the magnitude of eCO_2_-responsiveness. In contrast, infection with *P. sorghi* decreases the magnitude of eCO_2_-responsiveness across time. Infection with SCMV seems to have little impact on plant responses to eCO_2_.

**Fig. 6.**
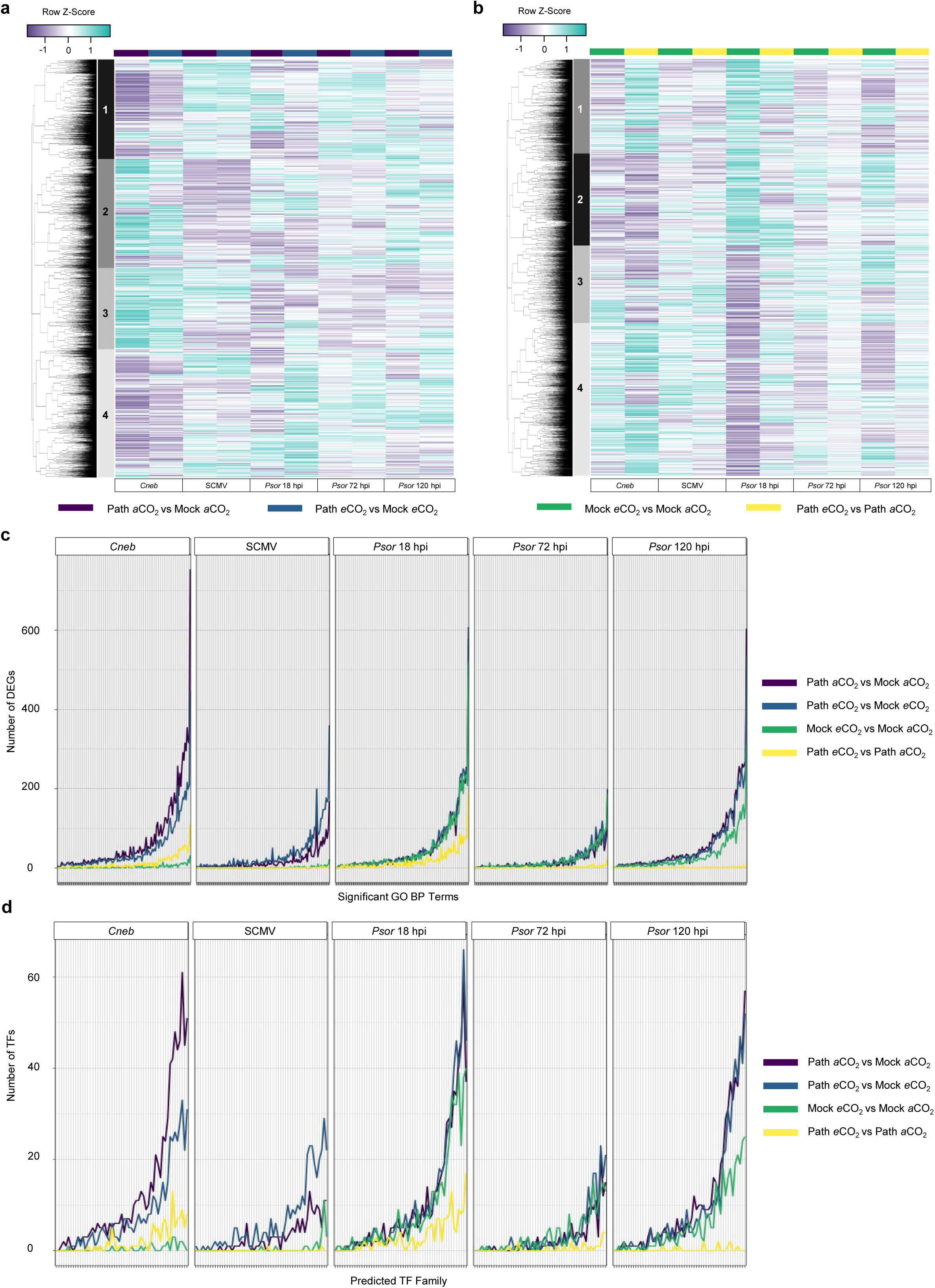
Identification of maize differentially expressed genes (DEGs), significantly overrepresented Gene Ontology (GO) terms, and predicted transcription factor (TF) families in response to pathogen infection and elevated CO_2_. **a**, Hierarchical clustering of 11,640 differentially expressed genes (DEGs; false discovery rate < 0.05) in maize leaves in response to three pathogen infections (*C. nebraskensis*, SCMV, and *P. sorghi*) compared with mock treatment under aCO_2_ (420 parts per million, ppm) or eCO_2_ (550 ppm). **b**, Reclustering of the same set of DEGs based on their transcriptional response to different CO_2_ levels across mock-treated and pathogen-inoculated samples. Samples for 3′ mRNA-Seq analysis were collected from the third leaves at 4 days post inoculation (dpi) with mock or *C. nebraskensis*, from the eleventh leaves at 14 dpi with mock or SCMV, and the from third leaves at 18, 72, and 120 hours post inoculation (hpi) with mock or *P. sorghi*. The three independent biological replicates (Rep) were conducted. Row Z-scores normalized log_2_ fold change values were used for hierarchical clustering of DEGs, based on expression across samples and replicates. Purple indicates expression values below the row mean for a given DEG and sample, and teal indicates expression values above the row mean for a given DEG and sample. **c**, Significantly overrepresented Gene Ontology (GO) term vs DEG number. **d**, Predicted transcription factor (TF) families vs TF number.

To better examine these responses, we returned to our pathogen-responsive (Path aCO_2_ vs Mock aCO_2_ and Path eCO_2_ vs Mock eCO_2_) and eCO_2_-responsive (Path eCO_2_ vs Path aCO_2_ and Mock eCO_2_ vs Mock aCO_2_) DEG lists (Supplementary Tables 17 and 18). Across all DEG lists, we identified 128 unique biological process terms significantly over-represented (corrected P < 0.05) in one or more DEG lists. For each pathogen and DEG list, we plotted the number of DEGs associated with each GO term (Fig. 6c). Examining pathogen-responsive DEGs, we identified more DEGs in aCO_2_ for *C. nebraskensis*, more DEGs in eCO_2_ for SCMV, and equal number of DEGs for *P. sorghi* across time points (Fig. 6c). Examining eCO_2_ responsive genes, we identified more DEGs in pathogen-inoculated samples for *C. nebraskensis* and few eCO_2_ responsive DEGs for SCMV. For *P. sorghi*, we see more eCO_2_ responsive DEGs in all three time points of mock-inoculated samples (Fig. 6c). Interestingly, eCO_2_ responsive DEGs seem to disappear at 72 and 120 hpi in *P. sorghi* inoculated samples. Also, worth noting is the eCO_2_ responsive DEGs in mock-inoculated samples at 18 and 72 hpi, closely mirror the pathogen-responsive DEGs under aCO*_2_* and eCO_2_ conditions. To see if this pattern holds true for DEGs with other functions, we queried our DEGs for known transcription factors (TFs) and plotted the number TFs relative to the 55 TF families. The same pattern observed in Fig 6c is repeated (Fig. 6d).

### Identification of regulatory elements governing responses to eCO_2_

To identify the TFs associated with eCO_2_ responses, we focused on the DEGs responding to [CO_2_] in pathogen- or mock-inoculated tissues (Path eCO_2_ vs Path aCO_2_ and Mock eCO_2_ vs Mock aCO_2_, FDR < 0.05, Supplementary Tables 6-15). Analyses were performed separately for *C. nebraskensis*, SCMV, and *P. sorghi*. Using PlantRegMap/PlantTFDB (version 5.0)^37^, we identified transcription factors (TFs) differently expressed in response to eCO_2_ and TF binding sites (TFBS) significantly overrepresented among the promoters of the DEGs in each list. To visualize the regulatory networks, the top Arabidopsis homolog for each TF was used as input to the STRING database (version 12)^38^.

For *C. nebraskensis* at 4 dpi, we identified 63 TFs/TFBS in mock plants (Fig. 7a) and 124 in infected plants (Fig. 7b) that were differentially expressed under eCO_2_ (Supplementary Table 19 and 20). These regulatory elements represented diverse families, including MYB, bZIP, WRKY, NAC, HD-ZIP, ERF, and bHLH, several of which are established regulators of plant stress and defense signaling. Infected plants contained more WRKY, NAC, HSF, and ARF family members, consistent with the activation of defense-related transcriptional programs under pathogen challenge compared with mock. A set of 53 TFs/TFBS was common to both conditions, indicating a conserved regulatory module responsive to eCO_2_. TFBS enrichment further revealed bZIP, MYB, NAC, Dof, TCP, and BBR-BPC motifs, with bZIP- and MYB-related elements consistently represented in both mock and infected plants, suggesting central roles in transcriptional regulation during *C. nebraskensis* infection in eCO_2_.

**Fig. 7.**
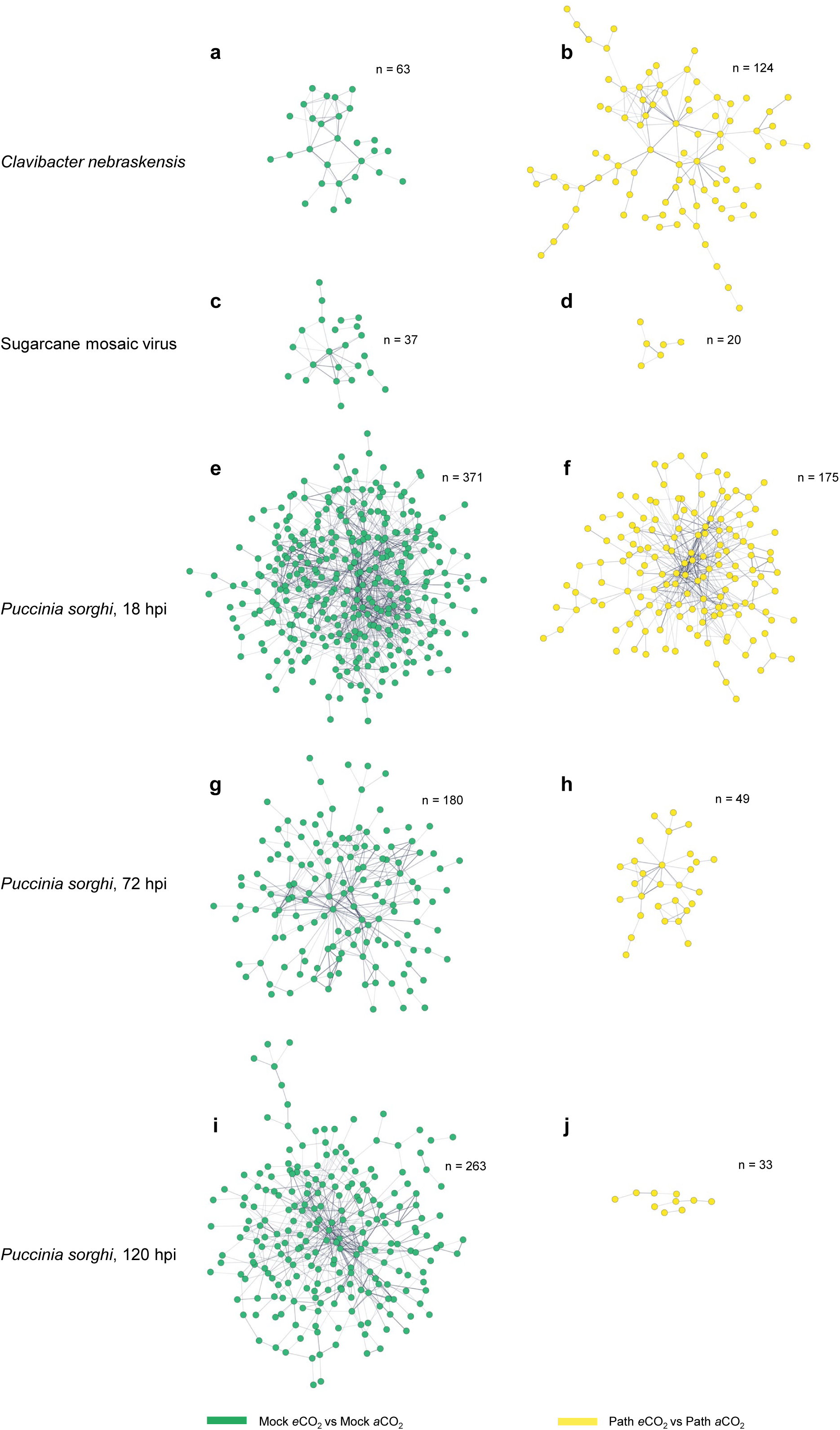
Transcription factor (TF) and transcription factor binding site (TFBS) networks associated with maize responses to elevated CO_2_ (eCO_2_) in mock-treated and pathogen-infected tissue. Differentially expressed genes (DEGs) responding to eCO_2_ in mock-treated or pathogen-infected tissues (Path eCO_2_ vs Path aCO_2_ and Mock eCO_2_ vs Mock aCO_2_, false discovery rate [FDR] < 0.05) were analyzed separately for *C. nebraskensis*, SCMV, and *P. sorghi*. TFs among DEGs were identified using PlantRegMap/PlantTFDB v5.0^37^, and significantly enriched TFBS were detected with the PlantRegMap enrichment tool. Arabidopsis homologs of maize TFs were mapped with a custom annotation tool and submitted to STRING v12 to infer functional association networks^38^. **a-b,** TF/TFBS networks for *C. nebraskensis*. **c-d,** TF/TFBS networks for SCMV. **e-f,** TF and TFBS networks for *P. sorghi* at 18 hpi. **g-h,** TF/TFBS networks for *P. sorghi* at 72 hpi. **i–j,** TF/TFBS networks for *P. sorghi* at 120 hpi. Nodes represent TFs or TFBS, edges indicate predicted associations, green nodes correspond to mock-treated tissue (Mock eCO_2_ vs Mock aCO_2_), and yellow nodes correspond to pathogen-inoculated tissue (Path eCO_2_ vs Path aCO_2_). *n* indicates the number of nodes in each network. Disconnected nodes were removed from the figure.

For SCMV at 14 dpi, 37 TFs/TFBS were identified in mock plants (Fig. 7c) and 20 in infected plants (Fig. 7d) in eCO_2_, with only two shared between conditions (Supplementary Tables 19 and 20). These regulators spanned multiple families, including WRKY, NAC, MYB, ERF, bHLH, HD-ZIP, TCP, GRAS, and MADS, several of which are associated with hormone signaling and stress-responsive transcriptional regulation. Compared with *C. nebraskensis*, the number of SCMV-responsive TFs/TFBS was much smaller, although differences in sampling time may also have contributed. TFBS enrichment highlighted ERF, MYB, bZIP, GATA, and TCP motifs, with ERF- and MYB-related elements consistently represented across conditions, suggesting potential roles in host transcriptional regulation during SCMV infection in eCO_2_.

For *P. sorghi*, analyses were conducted at three infection stages. At 18 hpi, 371 TFs/TFBS were identified in mock plants (Fig. 7e) and 175 in infected plants (Fig. 7f) under eCO_2_, with 147 shared between conditions (Supplementary Tables 19 and 20). At 72 hpi, 180 were identified in mock plants (Fig. 7g) and 49 in infected plants (Fig.7h), with 33 shared (Supplementary Tables 19 and 20). At 120 hpi, 263 were identified in mock plants (Fig. 7i) and 33 in infected plants (Fig. 7j), with 22 shared (Supplementary Tables 19 and 20). These regulators included families frequently associated with plant defense and hormone signaling, such as WRKY, NAC, MYB, ERF, bHLH, bZIP, HD-ZIP, ARF, TCP, GATA, and MADS. TFBS enrichment consistently highlighted bZIP, TCP, bHLH, HSF, MYB, Dof, and BBR-BPC motifs, with early infection showing the broadest representation and later stages displaying a more limited set. Together, these results suggest that *P. sorghi* infection under eCO_2_ initially engages a broad and diverse array of transcriptional regulators, which narrows as the infection progresses, potentially reflecting the combined influence of pathogen development and sampling time.

## Discussion

Rising atmospheric CO_2_ remains a defining challenge with profound implications for plant productivity and global food security^39^. In eCO_2_, plants exhibit altered physiological and biochemical processes that can potentially impact interactions with pathogenic microbes^17^. Although emerging research, primarily performed in C3 plants, sheds light on how eCO_2_ influences plant defense mechanisms, predicting the outcome of any particular plant-microbe interaction remains challenging^10^. Moreover, our understanding of how eCO_2_ affects C4 plant-pathogen interactions is essentially unknown. Our results demonstrate that predicted mid-century CO_2_ levels exert differential effects on maize immunity that vary across pathogen types.

The two prior studies that investigated the effects of eCO_2_ on a C4 plant-pathogen interaction specifically explored maize susceptibility to *Fusarium verticillioides* and its mycotoxin. Vaughan et al. (2014, 2016) found that eCO_2_ (800 ppm) increased maize susceptibility to *F. verticillioides* by compromising lipoxygenase-dependent signaling, leading to reduced jasmonic acid and phytoalexin levels, while fumonisin levels remained unchanged^23,24^. Under eCO_2_ conditions, diseases caused by biotrophic pathogens are often suppressed, whereas those caused by necrotrophs tend to be enhanced. Such variations likely stem from differences in plant species, cultivars, and ecotypes, as well as environmental factors such as [CO_2_], photoperiod, light intensity, and nutrients. We aimed to establish a systematic investigation to evaluate maize-pathogen responses under controlled environmental conditions to assess how eCO_2_ differentially affects different types of plant pathogens and to lay a foundation for understanding the complex and dynamic signaling events that influence maize-pathogen interactions in eCO_2_.

Several studies have addressed the impact of eCO_2_ on bacterial diseases in soybean^17^ infected with *Pseudomonas syringae* pv. *glycinea* (*Psg*) and in Arabidopsis^40,41^ and tomato^42–44^ infected with *P. syringae* pv. *tomato* DC3000 (*Pst*DC3000). Soybean grown under eCO_2_ (550 ppm) exhibited reduced susceptibility to *Psg*, consistent with enhanced immune responses^17^. Similarly, tomatoes grown under eCO_2_ (800 ppm) displayed decreased susceptibility to *Pst*DC3000, associated with enhanced SA-responsive gene expression and attenuation of JA-responsive gene expression^44^. In contrast, Arabidopsis grown under eCO_2_ (800 ppm) became more susceptible to *Pst*DC3000^40,41^. These discrepancies between Arabidopsis and tomato may reflect differences in species-specific immune regulation^40–45^. In our study, maize plants grown under eCO_2_ showed enhanced resistance to *C. nebraskensis*, as indicated by slower symptom progression (Fig. 2a), reduced lesion length (Fig. 2b,c), and lower bacterial proliferation (Fig. 2d,e). When assessing CO_2_ responsiveness, *C. nebraskensis*-infected plants displayed more eCO_2_-responsive defense regulators than mock plants, which may reflect the broader transcriptional activation occurring during infection (Fig. 7a,b), including members of the WRKY, NAC, and bZIP TF families. Taken together, it is apparent that eCO_2_ modulates bacterial immunity in both C3 and C4 plants in a pathosystem-specific manner.

PTI represents the first layer of inducible plant defense and is initiated when PRRs perceive MAMPs^46^. Both the study in soybean (C3)^17^ and our study in maize (C4) revealed several comparable PTI responses under eCO_2_, including increased ROS production (Fig. 2f), enhanced MAPK activation (Fig. 2h), and higher SA accumulation (Fig. 2j) or SA-marker gene (*GmPR1*) expression. In addition, maize plants grown under eCO_2_ displayed higher basal PA levels (Fig. 2g) accompanied by modest increases in RBOHD protein abundance (Fig. 2i), suggesting that eCO_2_ may enhance the biochemical conditions that support PTI signaling. These observations raise questions about whether early immune signaling components might intersect with the pathways plants use to perceive atmospheric [CO_2_]. Although CO_2_ perception is best studied in guard cells through the MAPK-HT1 regulatory module^47^, it is unknown whether similar sensing mechanisms function in other cell types. If CO_2_-responsive signaling extends beyond guard cells or involves additional receptors, changes in [CO_2_] may influence early PTI regulators and modify the strength of immune activation. From this perspective, eCO_2_ may shape early defense responsiveness by modulating signaling components shared between CO_2_ perception and innate immunity.

Studies in C3 plants demonstrate that eCO_2_ influences plant-virus interactions. For example, eCO_2_ enhances virus accumulation and disease severity for barley yellow dwarf virus (BYDV) in wheat^48,49^, tomato yellow leaf curl virus (TYLCV) in tomato^50^, and bean pod mottle virus and soybean mosaic virus in soybean^17^. In contrast, reduced virus titers and disease severity under eCO_2_ were reported for potato virus Y (PVY) in tobacco^51,52^, tobacco mosaic virus (TMV) in tomato^44^, and bean pod mottle virus in common bean^53^. Our results indicate that maize plants grown in eCO_2_ were more susceptible to SCMV infection, consistent with more limited transcriptional responses observed in eCO_2_ (Fig. 6c,d), where multiple *AGO1* and *AGO10* homologs were downregulated (Supplementary Table 8). Given the essential role of AGOs in RNA silencing, their downregulation suggests a plausible explanation for diminished antiviral immunity^54,55^. There are no studies that investigate roles of specific maize AGO proteins in antiviral defenses, so it will be interesting to test if these particular AGOs have significant antiviral roles in the future.

*P. sorghi* is an obligate biotroph that penetrates maize leaves through stomata, so changes in stomatal behavior under eCO_2_ could influence infection^56,57^. eCO_2_ was reported to alter disease development in rust fungi such as *P. striiformis* in cereals and *P. triticina* in wheat, potentially through changes in leaf morphology, stomatal behavior, and defense signaling^58,59^. As eCO_2_ generally reduces stomatal conductance and aperture in many plant species including maize (Fig. 4C), such changes could potentially influence spore germination or entry^60^. To determine whether these physiological changes extend to molecular responses, we assessed transcriptional responses. At the transcriptional level, *P. sorghi* infection induced a broad set of CO_2_-responsive regulators at 18 hpi, including WRKY, NAC, MYB, and bZIP families (Fig. 7e,f), with enrichment of immune- and stress-related motifs. This response was significantly repressed at 72 and 120 hpi, suggesting a temporal shift in host transcriptional activity as infection progressed (Fig. 7g-j). These early changes point to transient adjustments in defense regulation under eCO_2_, but they did not coincide with altered susceptibility, indicating that the infection outcome remained largely unchanged between CO_2_ treatments.

Our data indicate that maize plants are less susceptible to the hemibiotrophic *E. turcicum* and *C. graminicola* in eCO_2_, with no change in fungal growth *in vitro* (Supplementary Fig. 4b,5b). In barley, eCO_2_ (700 ppm) as a single factor increased basal resistance to powdery mildew (*Blumeria graminis* f. sp. *hordei*), although this benefit was not maintained when elevated temperature and ozone were combined with eCO_2_^61^. However, a three-year FACE study in rice reported higher susceptibility to blast (*Magnaporthe oryzae*) and a higher incidence of sheath blight (*Rhizoctonia solani*) under eCO_2_ (~580-660 ppm)^62^, demonstrating that eCO_2_ does not universally suppress fungal disease.

The effects of eCO_2_ on susceptibility to soil-borne pathogens are less studied than those of foliar pathogens^10^. eCO_2_ has been reported to induce changes in root architecture, exudate composition, and microbial community structure, which together can alter disease outcomes in a context-dependent manner^63^. For example, tomato plants grown under eCO_2_ (700 ppm) were more tolerant to root rot caused by *Phytophthora parasitica*^64^. In contrast, soybean plants grown under eCO_2_ (550 ppm) exhibited increased susceptibility to *Fusarium virguliforme*, as indicated by more severe sudden death syndrome (SDS) symptoms and higher fungal accumulation in roots, while their susceptibility to *Pythium sylvaticum* remained unchanged despite greater biomass loss^17^. Our data indicate that maize susceptibility to *P. sylvaticum* remains unchanged under eCO_2_; however, further research is needed to investigate how eCO_2_ alters root exudates and rhizosphere microbial communities, and how these changes directly affect maize interactions with soil-borne pathogens.

In conclusion, we demonstrate that eCO_2_ differentially alters maize defense responses and gene expression, leading to pathosystem-specific changes in susceptibility to bacterial, viral, and fungal pathogens. This study examined host physiology and disease development, but did not address how pathogen virulence, evolution, or host genetic diversity may further shape these interactions. An independent assessment of eCO_2_ was necessary for understanding its specific effects on plant immunity without the influence of confounding factors. However, other environmental variables such as temperature, water availability, and nutrient status are also expected to influence maize-pathogen dynamics. Understanding how these stresses interact with eCO_2_ will be critical for developing strategies to protect the health of maize and other C4 crops in the future.

## Methods

### Plant growth and maintenance

Sweet corn plants (*Zea mays* var Golden × Bantam; American Meadows) were used for all experiments due to their susceptibility to a wide array of pathogens, and the inbred H95 line was used for *Puccinia sorghi* infection assays. All plants were grown in CO_2_-controlled chambers at the Iowa State University Enviratron^65^, night (16°C and 70% relative humidity), day (28°C and 50% relative humidity), a 16-h light photoperiod, 420 ppm CO_2_ represented current aCO_2_ levels, and 550 ppm CO_2_ represented future eCO_2_ levels^66^. Plants were grown in LC-1 potting soil mix (Sungro horticulture, Catalog Number: 504307) and fertilized with 15-5-15 Cal-Mag Special (Peters Excel, G99140).

### Physiological measurements

In order to examine changes in maize physiology associated with eCO_2_, plant biomass, photosystem II (PSII) efficiency (quantum yield of fluorescence; ΦPSII efficiency), stomatal conductance (gas exchange rate; g_sw_), and transpiration (E) were measured for ten plants per chamber, with three chambers per CO_2_ treatment. Physiological measurements were conducted on the fully expanded fifth and sixth leaf between 21 and 42 days after planting (dap) using a LI-600 Porometer/Fluorometer (LI-COR Biosciences, Lincoln, NE, USA). Photosynthesis measurements were collected on the adaxial surface, while gas exchange measurements were simultaneously taken on the abaxial surface. At 42 dap, maize plants were assessed for height, measured from the stem base to the collar of the sixth leaf, and for shoot fresh weight, and were subsequently dried at 50°C for 3 days to determine shoot dry weight.

### Stomatal density and aperture measurements

Stomatal density and aperture measurements in the plant physiology experiment were obtained from ten leaf samples per CO_2_ chamber at 21 dap, with three chambers per CO_2_ treatment. For stomatal density measurements, impressions of the abaxial leaf surfaces were created using clear nail varnish^67^ and visualized using light microscopy (Leica DFC3000 G, 50× magnification; Leica Microsystems, Wetzlar, Germany). Stomatal aperture measurements for these samples were performed directly on the leaf tissue. Maize leaves were fixed in absolute ethanol for 1 h, washed with cold water, and submerged in a clearing solution (95% ethanol and 6–14% NaOCl in a 1:1 ratio, v/v) overnight. Prior to observation, leaf tissue was stained with 1 μM Rhodamine 6G (Sigma) for stomatal visualization^68^. For the SCMV and *P. sorghi* experiments, stomatal density and aperture were assessed using scanning electron microscopy. Impressions from both mock-treated and infected leaves were imaged using a NeoScope™ JCM-7000 Benchtop SEM (JEOL Ltd., Tokyo, Japan). Nail varnish impressions were placed directly into the low-vacuum chamber without sputter coating or ethanol dehydration, and three random fields of view per sample were counted. Stomatal aperture length, defined as the distance between the junctions or ends of the guard cells, was measured using ImageJ software^69^.

### RT-qPCR and qPCR analyses

Total RNA was extracted from approximately 50 mg of maize leaf tissue using TRIzol Reagent (Thermo Fisher Scientific, Waltham, MA, USA). Subsequently, 2 µg of RNA was reverse transcribed using the Maxima First Strand cDNA Synthesis Kit (Thermo Fisher Scientific, Waltham, MA, USA). RT-qPCR was conducted using the iTaq Universal SYBR Green Supermix kit (BioRad, Hercules, CA, USA) to quantify SCMV and assess the expression of *ZmPR1* (NM_001111929) and *ZmLOX9* (DQ335767). To quantify *P. sorghi*, *E. turcicum*, *C. graminicola*, and *P. sylvaticum*, total DNA was extracted from diseased tissues using the CTAB (cetyltrimethylammonium bromide) method^70^, and qPCR was performed using primers specific to each pathogen (Supplementary Table 1). ß-tubulin (NP_001105457) was used as an internal control for all experiments. New maize primers were designed based on the B73 reference genome version 5 (Zm-B73-REFERENCE-NAM-5.0; https://maizegdb.org) using the PrimerQuest Tool (Integrated DNA Technologies, Coralville, IA, USA).

### QuantSeq 3’ mRNA-Seq analyses

Total RNA was isolated from approximately 50 mg of maize leaf tissue using the TRIzol Reagent (Thermo Fisher Scientific, Waltham, MA, USA), followed by DNaseI treatment and purification using the RNA Clean and Concentrator-5 kit (Zymo Research, Irvine, CA, USA). RNA samples were quantified using the Qubit 4.0 Fluorometer (Invitrogen), and RNA integrity was evaluated using the Agilent 2100 Bioanalyzer (Iowa State University DNA Facility, Ames, IA). All RNA samples used for sequencing had an RQN (RNA Quality Number) equal to or > 6.5. Library preparation was performed using the QuantSeq 3’ mRNA-Seq Library Prep Kits (Lexogen, USA), and samples were sequenced using the Illumina Novaseq 6000 System (Iowa State University DNA Facility, Ames, Iowa).

For QuantSeq 3’ mRNA-Seq analyses, we generated 72 libraries for *C. nebraskensis* (2 pathogen treatments [mock or inoculated] x 6 chambers [3 aCO_2_ and 3 eCO_2_] x six plants per treatment combination), 60 libraries for SCMV (2 pathogen treatments x 5 chambers [2 aCO_2_ and 3 eCO_2_] x six plants per treatment combination), and 144 libraries for *P. sorghi* (2 pathogen treatments x 3 time points x] x 6 chambers [3 aCO_2_ and 3 eCO_2_] x four plants per treatment combination). Each library was sequenced on two lanes, generating two sequence files per library. FASTQC^71^ confirmed sequence quality and quantity. STAR version 2.7.10^72^ aligned reads to the Maize B73 reference genome (version Zm-B73-REFERENCE-NAM-5.0.fa), downloaded from MaizeGDB.org. Samtools version 1.17^73^ was used to merge the two mapping files (.bam) for each sample and identify uniquely mapping reads. For differential gene expression, samples from each pathogen were independently analyzed. Within RStudio version 4.4.1^74^, rtracklayer^75^ and summarizeOverlaps^75^ were used to assign reads from each sample to genes defined by the gene feature file corresponding to the reference genome. Sample replicates were visualized using ggplot2^76^ to ensure technical reproducibility. Following data visualization, 17 samples were removed from the *C. nebraskensis* analysis, 12 of which represented inoculated and mock-inoculated samples from a single eCO_2_ growth chamber. One sample was removed from each of the SCMV and *P. sorghi* analyses.

Following mapping and removal of samples, over 176.5, 260.4 and 759.8 million reads were used for differential gene expression analyses of *C. nebraskensis*, SCMV and *P. sorghi,* respectively. Trimmed Mean of M values in the Bioconductor package edgeR^77^ was used for data normalization. Only genes with log2 counts per million (cpm) > 1 in at least three replicates were used in the analysis. edgeR was used to identify differentially expressed genes (DEGs) among the different datasets using a false discovery rate of FDR < 0.05. For each pathogen, we identified DEGs responding to the pathogen (Pathogen-Mock) in eCO_2_ and aCO_2_ conditions. Similarly, we identified DEGs responding to eCO_2_ (eCO_2_-aCO_2_) in mock and pathogen-inoculated samples. For *P. sorghi*, this was repeated for each of the three time points. hclust^78^ was used to group similarly expressed DEGs of interest. Heatmaps were generated using Heatmap.2 in ggplot2^76^.

### DEG annotation and GO term enrichment

To annotate DEGs of interest, we developed a custom gene annotation tool for all predicted proteins in the Maize B73 reference genome (version Zm-B73-REFERENCE-NAM-5.0.fa). BLASTP^79^ was used to compare all predicted proteins against two custom versions of the Uniref100 databases^80^ (version 12/10/2024). In version 1, sequences with uninformative annotations were removed (hypothetical, uncharacterized, probable), while in version 2, sequences were taxonomically restricted to the 21 most represented plant species. In addition, BLASTP was used to compare predicted maize proteins against predicted proteins from the rice (Osa1_r7, version 03/02/25) and Arabidopsis (Araport 11, version 02/24/25) genomes. An E-value cutoff of 10^−10^ was used for all BLASTP comparisons. The top Osa1-R7 and Araport 11 hits were used to assign Gene Ontology (GO) terms for biological process, molecular function and cellular component^81^. The Arabidopsis GO terms were used to analyze GO term enrichment for DEG lists of interest using Fisher’s Exact Test and Bonferroni correction^82^. GO Terms were considered significantly over-represented with a corrected P-value < 0.05.

### Identification of transcription factors (TFs) and overrepresented transcription binding sites (TFBS)

To identify TFs among our DEGs, we took advantage of the Plant Transcriptional Regulatory Map/Plant Transcription Factor Database for *Z. mays*^37^ (PlantRegMap/PlantTFDB v5.0, https://plantregmap.gao-lab.org). Similarly, to identify over-represented TFBS, we used the PlantRegMap TF enrichment tool (https://plantregmap.gao-lab.org/tf_enrichment.php). To enable these analyses, DEGs were mapped to their version 3 names using a custom Perl script and a correspondence file provided by MaizeGDB (B73v3_B73v5_liftoff_genemodel_CDS_xref_shortened.txt, version 4/14/2025). Following these analyses, TF and TFs with overrepresented TFBS were mapped back to their v5 nomenclature. To visualize the network of TFs regulating responses to eCO_2_, their top Arabidopsis homologs, identified with the custom gene annotation tool above, were used as input to the STRING database (version 12)^38^.

### ROS burst assay

ROS burst assay was modified from a previous protocol for *Arabidopsis*^83^. Briefly, leaf discs were sampled from two-week-old plants grown in aCO_2_ or eCO_2_, placed in a 96-well plate containing 100 µL of water, and incubated overnight in the dark in CO_2_-controlled chambers. The next day, water was removed and replaced with 100 µL of elicitor solution containing 1 µM of the luminol derivative L-012 (Sigma-Aldrich), 10 µg/mL of horseradish peroxidase (HRP) (Sigma-Aldrich), and 1 µM of flg22 (VWR International, Radnor, PA, USA). Chemiluminescence was measured on a Synergy H1 microplate reader (BioTek Instruments, Winooski, VT, USA) every 2 min for 40 min with a 1 sec integration time.

### Plant protein extraction and immunoblot analysis

Total protein was extracted from maize leaves of two-week-old plants grown under aCO_2_ or eCO_2_. Tissues were ground in liquid nitrogen and resuspended at a 1:1 ratio (fresh weight/volume) in an extraction buffer, and protein concentrations were normalized using the Bradford assay (Thermo Fisher Scientific, Waltham, MA, USA)^84^. For detection of the RESPIRATORY BURST OXIDASE HOMOLOG D (RBOHD) protein accumulation, leaf tissues were treated with 1 µM flg22 peptide, and sampled from 0 to 30 min. RBOHD levels were assessed by immunoblotting using a primary anti-RBOHD antibody raised against Arabidopsis RBOHD (Agrisera, AS15 3002), which cross-reacts with maize RBOHDs, and a secondary goat anti-rabbit HRP-conjugated antibody (Cell Signaling, Danvers, MA, USA). For flg22-elicited MITOGEN-ACTIVATED PROTEIN KINASE (MAPK) activation assay, leaf tissues were treated with 1 µM flg22 peptide, and sampled from 0 to 120 min. MAPK activation was assessed by immunoblot analysis using a primary anti-Phospho-p44/42 MAPK antibody (Erk1/2; Thr-202/Tyr-204) (Cell Signaling, Danvers, MA, USA) and a secondary goat anti-rabbit-HRP antibody (Cell Signaling, Danvers, MA, USA).

### Preparation of samples for SA and JA LC-MS/MS quantification

A modified version of a methanolic extraction protocol established previously^17^ was used for sample preparation. Approximately 100 mg of each sample was ground and the extraction was initiated with the addition of 0.9 mL of 90% LC-MS grade methanol with 10% LC-MS grade water (Fisher Scientific, Waltham, MA) to each sample. Samples were also spiked with a standard addition of 10 µL of JA (2.5 µg/mL in ethyl acetate) and SA (2.5 µg/mL in 9:1 methanol:water) (Millipore Sigma, Burlington, MA), as well as 10 µL of deuterated SA-d4 (5 µg/mL in 9:1 methanol water) and JA-d5 (5 µg/mL in ethyl acetate) internal standards (Cayman Chemical, Ann Arbor, MI). Samples were then vortexed briefly and placed into an ice-cold sonication water bath for 10 min at full output power. Next, samples were vortexed for 5 min and centrifuged for 7 min at 15,000 xg. The supernatants were recovered, and the remaining insoluble pellets were re-extracted with an additional 0.9 mL of 90% methanol. The extracted supernatants were pooled.

An Oasis Hydrophilic-Lipophilic-Balanced (HLB) column (30 mg sorbent) (Waters Corporation, Milford, MA) was used for the purification of JA and SA. The column was first primed by washing with 1 mL of extraction solvent and then 0.5 mL of combined extract was loaded onto the column. The flow-through was collected and the column was washed with an additional 0.5 mL of extraction solvent. The sample was transferred to an LC vial and analyzed by LC-MS/MS. SA and JA standard curves with a range from 125 ng per sample down to 1.953 ng were prepared as serial dilutions in 9:1 methanol:water. The serial dilutions were treated in a corresponding manner as the tissue samples with the additions of extraction solvents along with the standard addition and internal standards, before being subjected to LC-MS/MS analysis.

### LC-MS/MS quantification of SA and JA

For LC-MS/MS SA and JA quantification, liquid chromatography separations were performed with an Agilent Technologies 1290 Infinity II UHPLC instrument equipped with an Agilent ZORBEX Eclipse plus C18 analytical column (2.1 mm × 100 mm, 1.8 µm) that was coupled 6470 triple quadrupole mass spectrometers Agilent Technologies, Santa Clara, CA). The LC-MS/MS analysis was performed using a modified version of the previously established conditions^85^. SA and JA sample extracts and standards were stored in the dark at 10°C in the autosampler during LC-MS/MS analysis. A volume of 6 µL of each sample was injected into the LC system. Chromatography was carried out at 40°C with a flow rate of 0.400 mL/min. Running solvents were A: water with 0.3 mM ammonium formate (pH adjusted to 3.5 using 0.1% formic acid) and B: acetonitrile with 0.3 mM ammonium formate (pH adjusted to 3.5 using 0.1% formic acid) (LC-MS grade, Fisher Scientific, Waltham, MA). Initial solvent conditions were 100% A which was held for 0.25-minutes which was then increased on a linear gradient to 65% B over 9.75-minutes, before further increasing to 100% B over a 2-minute gradient, 100% B was then held for 2-minutes before returning to 100% A over a 1-minute gradient. A 4-minute post run equilibration at 100% A was conducted after each LC-MS/MS acquisition.

Quantification analysis was performed in negative mode with the nozzle voltage set at 2000 V. Nitrogen was used as nebulizer, desolvation, and sheath gas. The nebulizer pressure was set at 40 psi with a capillary voltage of 2500 V. Desolvation gas was delivered at 12 L/min with temperature was heated to 350°C. The sheath gas flow was 10 L/min at a temperature of 350°C. High-purity nitrogen was used as a collision gas for collision induced dissociation. Multiple reaction monitoring (MRM) was used for detection with transitions of m/z 209→59 (JA), 214→62 (JA-d5), 137→93 (SA), and 141→97 (SA-d4), with all fragmentation conducted at 25 eV. All transitions were observed with a 100 ms dwell time while the cell accelerator and fragmentor were held at 5 V and 135 V respectively. Data evaluation and peak quantitation was performed using Agilent MassHunter Qualitative Analysis (version 10.0) and Agilent MassHunter Quantitative Analysis (version 10.0) software (Agilent Technologies, Santa Clara, CA). The SA and JA peaks were determined to have retention times of 5.15-minutes and 7.60-minutes respectively. While the peaks for the deuterated internal standards were found at 5.13-minutes (SA-d4) and 7.58-minutes. SA and JA quantification was finally determined by relative abundance to the internal standard and the SA and JA standard curve, before being made relative to the sample mass.

### Phosphatidic acid extraction

Approximately 100 mg of homogenized leaf tissue was used for sample preparation using modified versions of the Bligh and Dyer extraction and sample preparation methods^86,87^. Samples were spiked with internal standard, 10 µL of 15:0-18:1-d7-PA (1 mg/mL in chloroform (791642C, Avanti Polar Lipids, Birmingham, AL)). Extractions were initiated with the addition of 800 µL of 1:1 (v/v) of LC-MS grade methanol and 0.1 N HCl in LC-MS grade water (Fisher Scientific, Waltham, MA). Samples were vortexed for 1 minute and then placed in an ice-cold sonication water bath for 3 minutes. Next, 400 µL of reagent-grade chloroform (Fisher Scientific, Waltham, MA) was added, and the mixtures were vortexed for 3 minutes before centrifugation at 15,000 g for 7 minutes. The lower solvent layers were transferred to 0.2-µm centrifugal filters (Cat. No. F2517-9, Thermo Fisher Scientific, Waltham, MA). The extracts were then centrifuged for 3 minutes at 8,000 g. A PA standard curve (18:1 PA, 840875C, Avanti Polar Lipids, Birmingham, AL) was prepared in parallel with the sample extracts, with concentrations ranging from 19.5 to 5,000 ng per sample.

### Phosphatidic acid analysis via LC-MS/MS

Before LC-MS/MS analysis, sample extracts were filtered using 0.2-µm centrifugal filters (Cat. No. F2517-9, Thermo Fisher Scientific, Waltham, MA). LC separations were performed with an Agilent 1290 Infinity Binary Pump UHPLC system equipped with an Agilent Eclipse HILIC 1.8 µm, 2.1 × 100 mm analytical column, coupled to an Agilent 6540 UHD Accurate-Mass Q-TOF mass spectrometer (Agilent Technologies, Santa Clara, CA). Acquisition and data analysis were based on previously described methods^87^. A 2-µL volume of each sample was injected into the LC system. Chromatography was carried out at 40°C with a flow rate of 0.700 mL/min. The running solvents were: solvent A, water with 5 mM ammonium formate and 0.1% formic acid; and solvent B, 50% isopropanol in acetonitrile with 5 mM ammonium formate and 0.1% formic acid. Initial conditions were 100% B for 0.3 minutes, followed by a linear gradient to 65% B over 10 minutes. This was held for 5 minutes before returning to 100% B over a 2-minute gradient. A 6-minute post-run at 0% B was conducted after each LC-MS/MS acquisition.

Detection was performed using electrospray ionization in negative ion mode. Nitrogen was used as the source gas with a drying gas flow of 12 L/min at 350 °C, a nebulizer pressure of 32 psi, and a sheath gas flow of 11 L/min at 400 °C. The capillary and nozzle voltages were 4000 V and 1750 V, respectively. The mass spectrometer was operated in high-resolution (4 GHz) Auto MS/MS mode with a scan range of m/z 100–1700. The acquisition rate was 5 spectra/s for both MS and MS/MS scans, with a maximum of 4 MS/MS spectra acquired per cycle. Reference masses (m/z 112.985587 and 1033.988109) were continuously monitored for mass calibration during acquisition. Data evaluation and peak detection were performed using Agilent MassHunter Qualitative Analysis (version 10.0) and Mass Profiler (version 8.0). Peaks were identified by accurate mass spectral analysis in comparison with the METLIN Lipid database^88^ and the NIST20 spectral library (Agilent Technologies). Final quantification was performed with Agilent MassHunter Quantitative Analysis (version 10.0), calculated by integrating peak areas relative to the internal standard and standard curve, and normalized to sample mass.

### Bacterial infection and movement assays

*C. nebraskensis* was grown at 28°C overnight in nutrient broth-yeast extract medium (NBY)^89^. The next day, cells were harvested and resuspended in phosphate-buffered saline (PBS) solution (pH 7.0) and adjusted to OD_600_=0.4 (~10^8^ cells mL^−1^). The third leaf of two-week-old plants grown under aCO_2_ or eCO_2_ was inoculated with a bacterial suspension using a leaf-tip clipping method^90^. At 4 and 8 dpi, leaf discs were systematically extracted at distances of 5 and 10 cm from the inoculation site across six plants per CO_2_ treatment to quantify colony forming units (CFU/cm^2^) as described^91^. For 3′ mRNA-Seq analyses, samples were collected at 4 dpi between 09:00 h and 10:00 h from six mock-inoculated and six *C. nebraskensis*-infected plants per CO_2_ treatment in each chamber, yielding three biological replicates per treatment and 72 individual libraries in total (QuantSeq dataset 1).

### Bacterial growth curve

Liquid *C. nebraskensis* cultures were seeded to an initial OD_600_ of 0.02 in 100 mL of NBY broth. Cultures were grown in 500 mL Erlenmeyer flasks with gas-permeable caps to allow free diffusion of CO_2_ and were maintained in CO_2_-controlled growth chambers at either aCO_2_ or eCO_2_ at 26 °C with a rotation speed of 150 rpm^92^. Cultures were collected and monitored by recording OD_600_ every 4 h over 32 h.

### Viral infection assay

Fresh maize leaf tissues infected with SCMV were ground using a mortar in a 50 mM potassium phosphate buffer (pH 7.5). Approximately 10 mL of buffer was added to 1 g of tissues to make leaf sap. The seventh leaf of four-week-old plants was dusted with carborundum (Fisher Scientific, Waltham, MA) and mechanically inoculated by rubbing SCMV sap or buffer (mock). After rub-inoculations, the inoculated leaves were sprayed with water to remove excess carborundum dust. Plants were maintained under aCO_2_ or eCO_2_ as described above. For the assessment of viral titers, leaves of six plants were sampled, per treatment, at 14 and 21 dpi. Fresh weight and dry weight were measured at 21 dpi. Disease severity was evaluated between 0 and 21 dpi using a rating scale ranging from 0 to 3 (0 = no disease, 1 = mild, 2 = moderate, and 3 = severe). Area under disease progress curve (AUDPC) was calculated based on the equation shown below^93^. For 3′ mRNA-Seq analyses, leaf samples were collected at 14 dpi between 09:00 h and 10:00 h from six mock-treated and six SCMV-infected plants per CO_2_ treatment in each chamber, yielding three biological replicates per treatment and 60 individual libraries in total (QuantSeq dataset 2).

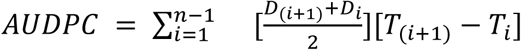

*D*_*i*_= disease severity on *i*th date, *T*_*i*_ = *i*th date, and *n* = total number of observations

### *P. sorghi* infection assays

For *P. sorghi* infection assays, 20 mg of frozen urediniospores of *P. sorghi* isolate IA16 were heat-activated for 5-10 min at 42°C and resuspended in 1 mL water with 1-2 drops of Tween 20 to create a spore suspension^94^. Urediniospore suspension was delivered onto the third and fourth leaves of 10-day-old susceptible H95 maize plants ^33^ grown at aCO_2_ or eCO_2_ using a procedure described by ^94^. The inoculated plants were misted with water and sealed in a plastic bag to create dew conditions and maintained at 20°C in the dark for 24 h^95^. For fungal biomass quantification, leaf tissues from two *P. sorghi*-infected maize plants were pooled per sample. For 3′ mRNA-Seq analyses, leaf tissues from two mock-inoculated or two *P. sorghi*-infected plants were pooled per sample at 18 hpi, 3 dpi, and 5 dpi, using eight plants per CO_2_ treatment in each chamber. This design yielded three biological replicates per treatment and 144 individual libraries in total (QuantSeq dataset 3). Leaves were imaged at 8 dpi, and the leaf pustule percentage coverage, mean pustule size, and number were calculated using a U-Net Neural Network trained for the *P. sorghi*-maize pathosystem^96^.

### *E. turcicum* infection assay

Sterilized millet was inoculated with a 14-day-old culture of *E. turcicum* and incubated at room temperature for 10 to 14 days^97^. Five mL of *E. turcicum*-infested millet was inoculated into the whorls of three-week-old maize plants maintained at aCO_2_ or eCO_2_. After inoculation, a high-humidity environment (80% relative humidity) was sustained for 3 days to promote the infection. At 28 dpi, leaves were collected from 6 plants, per CO_2_ treatment, to assess fungal titer. Disease severity was assessed using a 0 to 8 scale^98^ (Supplementary Table 2). AUDPC was calculated as described above.

### *C. graminicola* infection assay

*C. graminicola* was cultured on ½ strength PDA at 25°C for 7 days. Spores were harvested by flooding the plate with 0.02% Tween 20 (v/v) in sterile water and filtering through a sterilized cheesecloth. The final concentration was adjusted to ~10^6^ spores/mL^99^. Leaf sheaths of six-week-old plants grown at aCO_2_ or eCO_2_ were removed to expose the second internodes. The second internode was then punctured with a sterile scalpel blade, and a toothpick soaked in the spore suspension was inserted to 1/3 of its length (2.7 cm) carrying approximately 0.03 mL of suspension (~30,000 spores). At 28 dpi, stalk tissues were sampled from 6 plants of each CO_2_ treatment. Disease severity was assessed on a 0 to 5 rating scale as described^100^ (Supplementary Table 3).

### *E. turcicum* and *C. graminicola* plate assays

Starting cultures of *E. turcicum* and *C. graminicola* were grown on one-half strength PDA at room temperature for 14 days, respectively. A 7-mm plug was punched out of the starting culture and placed into the center of a 90-mm-diameter Petri dish containing ½ strength PDA. Ten plates, per CO_2_ treatment, were maintained under aCO_2_ or eCO_2_ ppm. Fungal growth was assessed every other day for 15 days.

### *P. sylvaticum* infection assay

Millet was inoculated with a 3-day-old culture of *P. sylvaticum* (isolate Gr8) grown on 4% V8 juice media containing neomycin sulfate (50 µg/ml) and chloramphenicol (10 µg/ml) (DV8++). The millet was colonized in the absence of light at room temperature for 7 to 14 days before use. For assay preparation, a mixture of 100 mL of sterile sand soil and 10 mL of *P. sylvaticum*-infested millet was initially placed in a 237 mL (8 oz.) polystyrene cup and covered with 25 mL of peat mix. The seed was positioned on top of the peat mix to be separated from the inoculum, and an additional layer of 25 mL of peat mix was added on top of the seed. The cups were incubated under aCO_2_ or eCO_2_, as described above. At 14 dap, 6 cups from each treatment were destructively sampled. Disease severity for each seedling was assessed using the 0 to 4 rating scale^101^ (Supplementary Table 4). In addition, fresh and dry weights of shoots and roots were recorded.

### Seed rot assay

A modified pathogenicity test^101,102^ was used to assess seed rot caused by *P. sylvaticum.* Cultures of *P. sylvaticum* were grown on 9-cm Petri plates with DV8++ in the dark under aCO_2_ or eCO_2_ for 3 days, until mycelia colonized the entire plate. Seeds were surface sterilized in a 1% sodium hypochlorite solution for 3 min, followed by a 3-min rinse in sterile water, and drying on sterilized paper towels. Five seeds were equidistantly spaced on a *P. sylvaticum*-inoculated plate. To evaluate germination as part of seed quality control, 10 seeds were placed on a non-inoculated DV8++ plate. The plates were incubated under aCO_2_ or eCO_2_ for 7 days. To assess seed rot, a modified 0-4 rating scale was used^101^ (Supplementary Table 5).

### Statistical analysis

Linear mixed-effects models (LMMs) were fit to each response according to the experimental design. We use the bacterial infection experiment measuring colony-forming units (CFUs) as an example to illustrate our LMM analysis. The experimental design is as follows. At the whole-plot level, growth chambers were randomly assigned to the two CO_2_ levels. Within each chamber, rows of plants (split-plots) were randomly selected to be destructively harvested at each dpi (days post-inoculation). For each individual plant, CFUs were measured at two distances from the inoculation site. Following this design, the fixed effects of the LMM included the main effects and all interaction effects of the factors CO_2_, dpi, and distance, while the random effects accounted for chamber, split-plot, and plants. Similar to this example, LMMs were fit to other responses according to the corresponding experimental designs.

For response variables that exhibited unequal variances on the original scale, LMMs were fit to log-transformed data, and standard errors (SE) on the original scale were obtained using the Delta method. For all LMM analyses, model diagnostic checks were performed to ensure that model assumptions were appropriate. Model parameters were estimated using the lmer() function in the lme4 package. The emmeans() function in the emmeans package was used to compute SEs. Type III ANOVA tables with F-tests were generated using the anova() function in combination with the lmertest package.

For response variables that exhibited unequal variances even after log-transformation, the linear mixed effect model with unequal variance was fitted on the original data. Model parameters were estimated using glmmTMB() function in glmmTMB R package. The emmeans() function in the emmeans package was used to compute SEs. Significance of fixed effects was determined by Type III Wald Chi-square tests using Anova() function in the car R package.

For ordinal response variables like disease score and root rot assay, cumulative link mixed model (CLMM) was fit for each response according to the experimental design. The functions clmm() or clmm2() in the ordinal package were used to estimate model parameters estimation and conduct Chi-square tests for fixed effects.

## Data availability

The RNA-seq reads for QuantSeq 3’ mRNA-Seq datasets 1, 2, and 3 were deposited in NCBI under BioProject PRJNA1206457, PRJNA1206458, and PRJNA1206459, respectively. The raw data and R-generated statistical reports supporting Fig. 1-5 and Supplementary Fig. 1-6 were deposited at the Iowa State University DataShare, and they can be accessed using this DOI: 10.25380/iastate.30875303.

## Acknowledgments

We are grateful to Dr. Alison Robertson (ISU) and Dr. Clarice Schmidt (ISU) for providing advice on maize pathogens, guidance on inoculation methods and infection assays, and cultures of Iowa isolates of *Clavibacter nebraskensis*, *Pythium sylvaticum*, *Exserohilum turcicum*, and *Colletotrichum graminicola*. We thank the ISU DNA Facility and the W. M. Keck Metabolomics Research Laboratory (RRID:SCR_017911) for providing analytical instrumentation and technical expertise. We thank Dr. Katerina Holan (ISU) for assistance with automated scoring of common rust disease. This work was supported by the ISU Plant Sciences Institute and USDA NIFA Hatch Project IOW04308. Ekkachai Khwanbua was supported by Anandamahidol fellowship from the government of Thailand. This research was funded in part by USDA-ARS project 5030-21220-007-000D and used resources provided by the USDA SCINet, USDA-ARS project 0500-00093-001-00-D. Mention of trade names or commercial products in this publication is solely for the purpose of providing specific information and does not imply recommendation or endorsement by the U.S. Department of Agriculture. USDA is an equal opportunity employer and provider.

## Contributions

EK and SAW conceived and designed the study. EK performed most of the experiments. JS assisted with the *Puccinia sorghi* (common rust) experiment. YQ and PL conducted the statistical analyses using linear mixed models. MAG performed the 3’ mRNA QuantSeq data analysis. EK prepared the initial manuscript draft with input from all authors. SAW and MAG provided review and editing, and all authors approved the final version of the manuscript.

## Ethics declaration

### Supplementary information

#### Source data

##### Figure legend (Supplementary)

**Supplementary Fig. 1 Effects of elevated CO_2_ on the growth rate of *Clavibacter nebraskensis*.** Bacterial cultures were grown in Nutrient Broth Yeast (NBY) without antibiotics for 32 h, and optical density at 600 nm (OD_600_) was recorded every 4 h using a spectrophotometer. Three independent experimental replicates were performed simultaneously for each CO_2_ concentration, in independent CO_2_-controlled chambers. Data points represent the average OD_600_ across replicates at each time point. *P*-value for the bacterial growth curves was obtained from the *F*-test of the main CO_2_ effect on log-transformed data in a linear mixed-effects model, and means with SE are presented on the log10 scale. The letter e denotes an exponent to the power of 10, and ppm denotes parts per million.

**Supplementary Fig. 2 Activation of MAPKs and accumulation of RBOH protein in maize leaves following flg22 treatment under ambient or elevated CO_2_.** MAPK activation (**a-b**) and RBOHD accumulation (**c,d**) were analyzed by immunoblot using maize leaves collected at the indicated time points after flg22 treatment under ambient or elevated CO_2_. Six plants per treatment were sampled at each CO_2_ concentration. MAPK activation was detected with anti-p42/44 MAPK antibody using protein extracts collected at 0, 10, 30, 60, 90, and 120 min. RBOH accumulation was detected with anti-AtRBOHD antibody using leaf tissues collected at 0, 15, and 30 min, with Coomassie Brilliant Blue (CBB) staining as the loading control. These blots represent two additional independent replicates in addition to the replicate shown in Fig. 2.

**Supplementary Fig. 3 Effects of elevated CO_2_ on shoot biomass and physiological traits in mock- and sugarcane mosaic virus (SCMV)-inoculated maize plants and on SCMV accumulation at 21 days post inoculation (dpi). a-b**, Height to the 9^th^ leaf collar and shoot fresh weight of maize plants at 21 dpi with SCMV or mock treatment. **c**, SCMV accumulation at 21 dpi was measured using reverse transcriptase-quantitative PCR (RT-qPCR) with primers specific to SCMV. Absolute quantification was performed using a standard curve generated from serial dilutions of the SCMV infectious clone. **d-f**, Stomatal conductance (g_sw_) and transpiration rate (E), and photosystem II efficiency (ΦPSII) was measured on the eleventh leaf of mock- and SCMV-treated maize plants at 21 dpi using the LI-600 system. Three experiments were conducted simultaneously in independent CO_2_ control chambers, using 6 plants per treatment for each CO_2_ concentration according to a replicated complete design. Data points represent the mean value across the three replicates and error bars represent the standard error. Due to unequal variance on the original scale, linear mixed effect model was applied to the log-transformed SCMV quantity and E data, and means with SE are presented on the log10 scale. *P*-values for plant height and shoot fresh weight were obtained from *F*-tests of the interaction effect. The *P*-value for SCMV quantity was obtained from the pairwise *t*-test comparing the two CO_2_ levels at 21 dpi. *P*-values for g_sw_, E, and ΦPSII were obtained from *t*-tests of the interaction effect at 21dpi. The letter e denotes an exponent to the power of 10, and ppm denotes parts per million.

**Supplementary Fig. 4 Relative expression of the JA marker gene *ZmLOX9* and *in vitro* growth of *Exserohilum turcicum* under eCO_2_. a**, Relative expression of *ZmLOX9* (JA marker gene) in maize was assessed by reverse transcriptase quantitative polymerase chain reaction (RT-qPCR) analysis relative to the *Zmβ-TUB*. Three experiments were conducted simultaneously in independent CO_2_ control chambers, using 6 plants per treatment for each CO_2_ concentration according to a replicated complete randomized design. **b**, Hyphae diameter of *E. turcicum* was measured every other day over 15 days. Three experiments were conducted simultaneously using 10 plates per experiment for each CO_2_ treatment. Data were graphed as the mean across the three replicates with SE bars. *P*-value for *ZmLOX9* expression was obtained from *F*-tests of the interaction effect from the linear mixed effect model (LMM) analysis of the log-transformed data to account for unequal variances, and means with SE are presented on the log10 scale. *P*-value for the plate assay was obtained from *F*-tests of the main CO_2_ effect from the LMM analysis. The letter e denotes an exponent to the power of 10, and ppm denotes parts per million.

**Supplementary Fig. 5 Relative expression of the JA marker gene *ZmLOX9* and *in vitro* growth of *Colletotrichum graminicola* under eCO_2_. a**, Relative expression of *ZmLOX9* (JA marker gene) in maize was quantified by reverse transcriptase quantitative polymerase chain reaction (RT-qPCR) analysis relative to the *Zmβ-TUB*. Three experiments were conducted simultaneously in independent CO_2_ control chambers, using 6 plants per treatment for each CO_2_ concentration according to a replicated complete randomized design. **b**, Hyphae diameter of *C. graminicola* was measured every other day over 15 days. Three experiments were conducted simultaneously using 10 plates per experiment, for each CO_2_ treatment. Data were graphed as the mean across the three replicates with SE bars. *P*-value for *ZmLOX9* expression was obtained from *F*-tests of the main CO_2_ effect from the linear mixed effect model (LMM) analysis of the log-transformed data to account for unequal variances, and means with SE are presented on the log10 scale. *P*-value for the plate assay was obtained from *F*-tests of the main CO_2_ effect from the LMM analysis. The letter e denotes an exponent to the power of 10, and ppm denotes parts per million.

**Supplementary Fig. 6 Effects of elevated CO_2_ on shoot and root biomass, *ZmPR1* expression, and seed decay in maize infected with *Pythium sylvaticum*. a-c**, Shoot fresh weight, root fresh weight, and shoot dry weight of maize plants at 14 dpi with *P. sylvaticum* or mock treatment. **d**, Relative expression of *ZmPR1*, a salicylic acid (SA) marker gene, was assessed by RT-qPCR relative to *Zmβ-TUB*. Three experiments were conducted simultaneously in independent CO_2_ control chambers, using 6 plants per treatment for each CO_2_ concentration according to a replicated complete randomized design. **e**, Seed rot severity was evaluated using a 0-4 scale, where 0 indicated that the seed germinated with no radicle discoloration; 1 indicated germination with less than 50% radicle discoloration; 2 indicated germination with more than 50% radicle discoloration; 3 indicated that the seed germinated but subsequently rotted; and 4 indicated that the seed rotted without germinating. Five maize seeds were placed equidistantly on *P. sylvaticum*-inoculated plates. Ten plates per CO_2_ treatment were incubated for 7 days under either ambient (aCO_2_) or elevated (eCO_2_) CO_2_ conditions. *P*-values for shoot fresh weight, root fresh weight, shoot dry weight, and *ZmPR1* expression were obtained from *F*-tests of the interaction effect from the linear mixed effect model analysis of the log-transformed data to account for unequal variances, and means with SE are presented on the log10 scale. *P*-value for the seed-rot assay was obtained from a likelihood ratio test in a cumulative link mixed model. The letter e denotes an exponent to the power of 10, and ppm denotes parts per million.

**Supplementary Fig. 7 Hierarchical clustering of differentially expressed genes (DEGs; false discovery rate < 0.05) in maize leaves at 4 days post inoculation (dpi) with mock or *Clavibacter nebraskensis* under ambient (aCO_2_, 420 parts per million (ppm)) or elevated CO_2_ (eCO_2_, 550 ppm). a**, Clustering of 14,793 DEGs responding to *C. nebraskensis* at 4 dpi in plants grown in aCO_2_ or eCO_2_. Four distinct expression clusters were identified and are indicated by differently shaded gray boxes to the left of the heat map. The full list of DEGs is provided in Supplementary Table 6. **b**, Clustering of 2,498 DEGs responding to eCO_2_ (420 ppm vs 550 ppm) in either mock or *C. nebraskensis*-infected samples at 4 dpi. Three different expression clusters were identified and are indicated by differently shaded gray boxes to the left of the heat map. The full list of DEGs is provided in Supplementary Table 7. Rep indicates independent biological replicates. For each CO_2_ condition, three replicates were performed for mock and *Clavibacter nebraskensis* treatments under aCO_2_, and two replicates were performed under eCO_2_. Each replicate included six plants. Row Z-scores were used for hierarchical clustering of DEGs, based on expression across samples and replicates. Purple indicates expression values below the row mean and teal indicates expression values above the row mean.

**Supplementary Fig. 8 Hierarchical clustering of differentially expressed genes (DEGs; false discovery rate < 0.05) in maize leaves at 14 days post inoculation (dpi) with mock or sugarcane mosaic virus (SCMV) under ambient (aCO_2_, 420 parts per million (ppm)) or elevated CO_2_ (eCO_2_, 550 ppm). a**, Clustering of 4,210 DEGs responsive to SCMV at 14 dpi in plants grown in aCO_2_ or eCO_2_. Two distinct expression clusters were identified and are indicated by differently shaded gray boxes to the left of the heat map. The full list of DEGs is provided in Supplementary Table 8. **b**, Clustering of 176 DEGs responding to eCO_2_ (420 ppm vs 550 ppm) in either mock or SCMV-infected samples at 14 dpi. Three different expression clusters were identified and are indicated by differently shaded gray boxes to the left of the heat map. The full list of DEGs is provided in Supplementary Table 9. Rep indicates independent biological replicates. For each CO_2_ condition, two replicates were performed for mock and SCMV treatments under aCO_2_, and three replicates were performed under eCO_2_. Row Z-scores were used for hierarchical clustering of DEGs, based on expression across samples and replicates. Purple indicates expression values below the row mean and teal indicates expression values above the row mean.

**Supplementary Fig. 9 Hierarchical clustering of differentially expressed genes (DEGs; false discovery rate < 0.05) in maize leaves at 18 hours post inoculation (hpi) with mock or *Puccinia sorghi* under ambient (aCO_2_, 420 parts per million (ppm)) or elevated CO_2_ (eCO_2_, 550 ppm). a**, Clustering of 9,671 DEGs responsive to *P. sorghi* at 18 hpi in plants grown in aCO_2_ or eCO_2_. Two distinct expression clusters were identified and are indicated by differently shaded gray boxes to the left of the heat map. The full list of DEGs is provided in Supplementary Table 10. **b**, Clustering of 10,189 DEGs responding to eCO_2_ (420 ppm vs 550 ppm) in either mock or *P. sorghi*-infected samples at 18 hpi. Three different expression clusters were identified and are indicated by differently shaded gray boxes to the left of the heat map. The full list of DEGs is provided in Supplementary Table 11. Rep indicates independent biological replicates. The three independent replicates were conducted using eight plants per CO_2_ treatment. Row Z-scores were used for hierarchical clustering of DEGs, based on expression across samples and replicates. Purple indicates expression values below the row mean and teal indicates expression values above the row mean.

**Supplementary Fig. 10 Hierarchical clustering of differentially expressed genes (DEGs; false discovery rate < 0.05) in maize leaves at 72 hours post inoculation (hpi) with mock or *Puccinia sorghi* under ambient (aCO_2_, 420 parts per million (ppm)) or elevated CO_2_ (eCO_2_, 550 ppm). a**, Clustering of 5,035 DEGs responsive to *P. sorghi* at 72 hpi in plants grown in aCO_2_ or eCO_2_. Two distinct expression clusters were identified and are indicated by differently shaded gray boxes to the left of the heat map. The full list of DEGs is provided in Supplementary Table 12. **b**, Clustering of 3,674 DEGs responding to eCO_2_ (420 ppm vs 550 ppm) in either mock or *P. sorghi*-infected samples at 72 hpi. Two different expression clusters were identified and are indicated by differently shaded gray boxes to the left of the heat map. The full list of DEGs is provided in Supplementary Table 13. Rep indicates independent biological replicates. The three independent replicates were conducted using eight plants per CO_2_ treatment. Row Z-scores were used for hierarchical clustering of DEGs, based on expression across samples and replicates. Purple indicates expression values below the row mean and teal indicates expression values above the row mean.

**Supplementary Fig. 11 Hierarchical clustering of differentially expressed genes (DEGs; false discovery rate < 0.05) in maize leaves at 120 hours post inoculation (hpi) with mock or *Puccinia sorghi* under ambient (aCO_2_, 420 parts per million (ppm)) or elevated CO_2_ (eCO_2_, 550 ppm). a**, Clustering of 13,494 DEGs responsive to *P. sorghi* at 120 hpi in plants grown in aCO_2_ or eCO_2_. Three distinct expression clusters were identified and are indicated by differently shaded gray boxes to the left of the heat map. The full list of DEGs is provided in Supplementary Table 14. **b**, Clustering of 5,624 DEGs responding to eCO_2_ (420 ppm vs 550 ppm) in either mock or *P. sorghi*-infected samples at 120 hpi. Six different expression clusters were identified and are indicated by differently shaded gray boxes to the left of the heat map. The full list of DEGs is provided in Supplementary Table 15. Rep indicates independent biological replicates. The three independent replicates were conducted using eight plants per CO_2_ treatment. Row Z-scores were used for hierarchical clustering of DEGs, based on expression across samples and replicates. Purple indicates expression values below the row mean and teal indicates expression values above the row mean.

##### Supplementary tables

**Supplementary Table 1.**
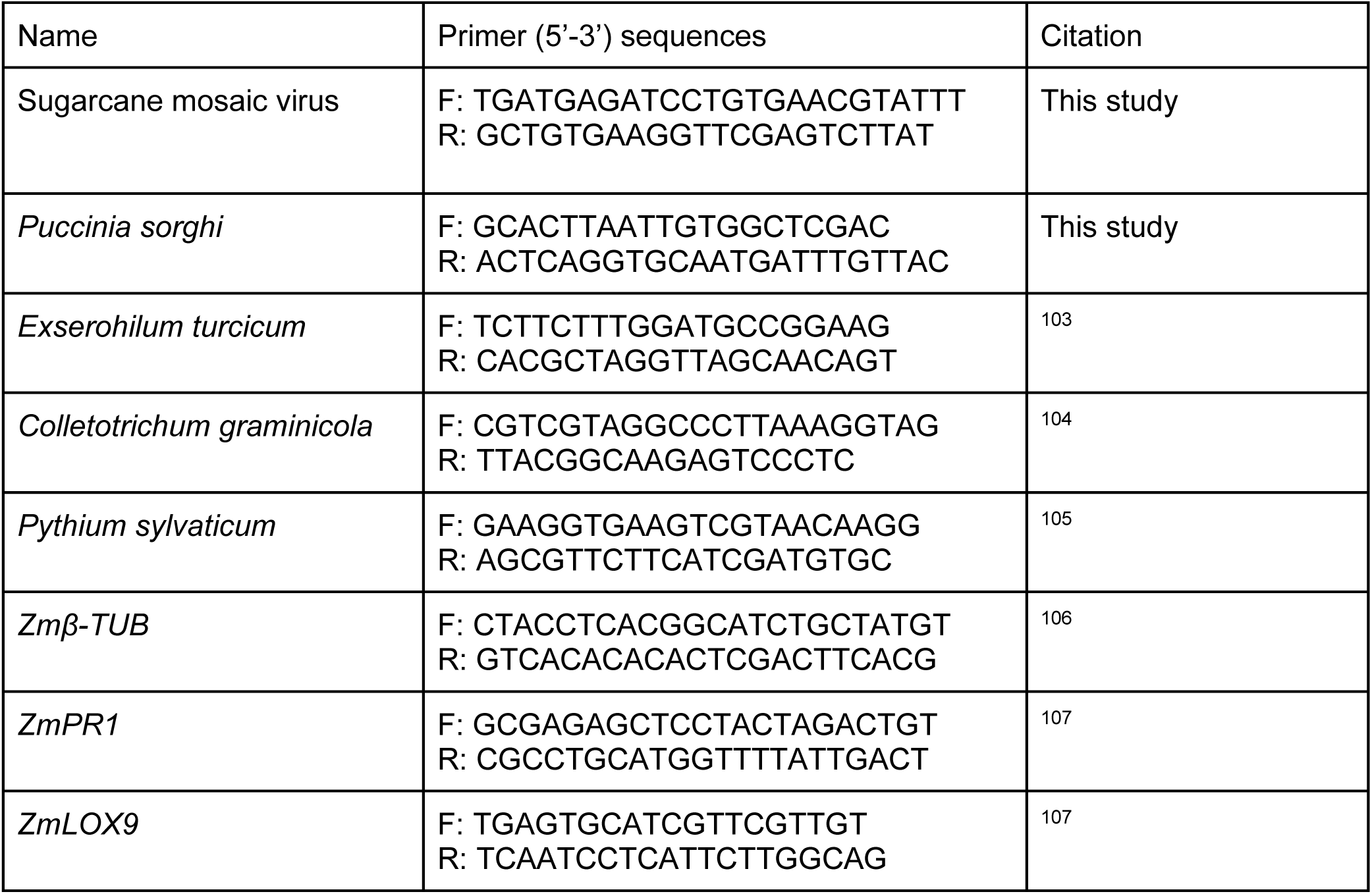
Oligonucleotide primers used in this study.

**Supplementary Table 2.**
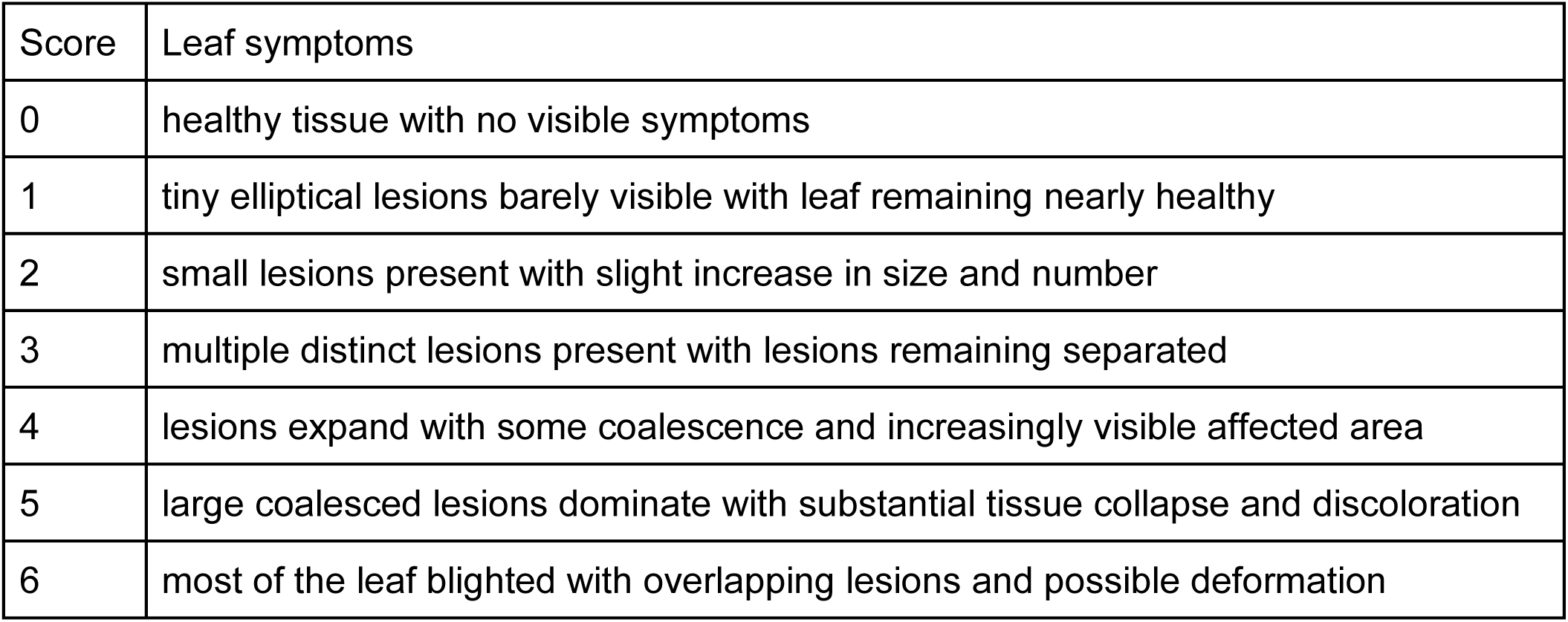
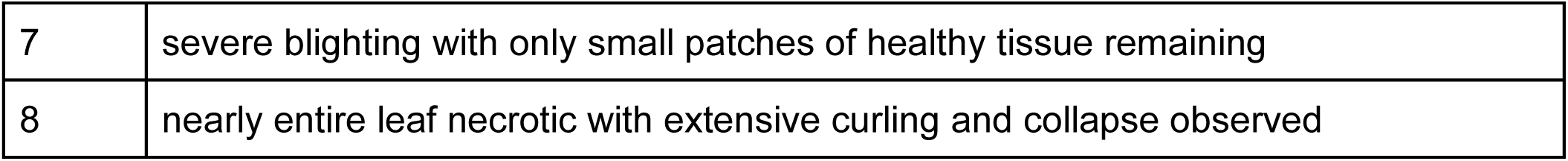
Disease rating scale of Northern corn leaf blight ranging from 0 to 898.

**Supplementary Table 3.** Disease rating scale of stalk rot severity ranging from 0 to 5100.

| Score | Stalk symptoms |
| --- | --- |
| 0 | no visible symptoms; pith remains white, firm, and intact |
| 1 | small localized lesions with minor discoloration |
| 2 | visible patches of necrosis covering ~25% of the tissue; some pith degradation |
| 3 | ~50% of the stalk necrotic; clear tissue collapse and dark discoloration |
| 4 | extensive necrosis (~75%); widespread tissue degradation |
| 5 | entire stalk internode necrotic; fully degraded pith with loss of structural integrity |

**Supplementary Table 4.** Disease rating scale of root symptoms ranging for 0 to 4101.

| Score | Root symptoms |
| --- | --- |
| 0 | root were long and full with no signs of discoloration or infection by the pathogen |
| 1 | roots were slightly stunted and moderately discolored |
| 2 | roots were severely stunted and may be severely discolored |
| 3 | roots were very short and rotted |
| 4 | seed was rotted and had not germinated |

**Supplementary Table 5.** Seed rot severity scale ranging from 0 to 4101.

| Score | Seed rot severity |
| --- | --- |
| 0 | seed germinated and there was no radicle discoloration |
| 1 | seed germinated and < 50% of the radicle was discolored |
| 2 | seed germinated but > 50% of the radicle was discolored |
| 3 | seed had germinated but then rotted |
| 4 | seed rotted without germinating |

**Supplementary Table 6. Significantly differentially expressed genes (DEG, false discovery rate (FDR) < 0.05) responding to *Clavibacter nebraskensis* (*Cneb*) infection at 4 days post-inoculation (dpi) in plants grown at elevated CO_2_ (eCO_2_, 550 ppm) or ambient CO_2_ (aCO_2_, 420 ppm) conditions.**

**Supplementary Table 7. Significantly differentially expressed genes (DEG, false discovery rate (FDR) < 0.05) responding to changes in CO_2_ at elevated CO_2_ (eCO_2_, 550 ppm) or ambient CO_2_ (aCO_2_, 420 ppm) in *Clavibacter nebraskensis* (*Cneb*)-infected plants or mock-inoculated plants at 4 days post-inoculation (dpi).**

**Supplementary Table 8. Significantly differentially expressed genes (DEG, false discovery rate (FDR) < 0.05) responding to sugarcane mosaic virus (SCMV) infection at 14 days post-inoculation (dpi) in plants grown at elevated CO_2_ (eCO_2_, 550 ppm) or ambient CO_2_ (aCO_2_, 420 ppm) conditions.**

**Supplementary Table 9. Significantly differentially expressed genes (DEG, false discovery rate (FDR) < 0.05) responding to changes in CO_2_ at elevated CO_2_ (eCO_2_, 550 ppm) or ambient CO_2_ (aCO_2_, 420 ppm) in sugarcane mosaic virus (SCMV)-infected plants or mock-inoculated plants at 14 days post-inoculation (dpi).**

**Supplementary Table 10. Significantly differentially expressed genes (DEG, false discovery rate (FDR) < 0.05) responding to *Puccinia sorghi* (*Psor*) infection at 18 hours post-inoculation (hpi) in plants grown at elevated CO_2_ (eCO_2_, 550 ppm) or ambient CO_2_ (aCO_2_, 420 ppm) conditions.**

**Supplementary Table 11. Significantly differentially expressed genes (DEG, false discovery rate (FDR) < 0.05) responding to changes in CO_2_ at elevated CO_2_ (eCO_2_, 550 ppm) or ambient CO_2_ (aCO_2_, 420 ppm) in *Puccinia sorghi* (*Psor*)-infected plants or mock-inoculated plants at 18 hours post-inoculation (hpi).**

**Supplementary Table 12. Significantly differentially expressed genes (DEG, false discovery rate (FDR) < 0.05) responding to *Puccinia sorghi* (*Psor*) infection at 72 hours post-inoculation (hpi) in plants grown at elevated CO_2_ (eCO_2_, 550 ppm) or ambient CO_2_ (aCO_2_, 420 ppm) conditions.**

**Supplementary Table 13. Significantly differentially expressed genes (DEG, false discovery rate (FDR) < 0.05) responding to changes in CO_2_ at elevated CO_2_ (eCO_2_, 550 ppm) or ambient CO_2_ (aCO_2_, 420 ppm) in *Puccinia sorghi* (*Psor*)-infected plants or mock-inoculated plants at 72 hours post-inoculation (hpi).**

**Supplementary Table 14. Significantly differentially expressed genes (DEG, false discovery rate (FDR) < 0.05) responding to *Puccinia sorghi* (*Psor*) infection at 120 hours post-inoculation (hpi) in plants grown at elevated CO_2_ (eCO_2_, 550 ppm) or ambient CO_2_ (aCO_2_, 420 ppm) conditions.**

**Supplementary Table 15. Significantly differentially expressed genes (DEG, false discovery rate (FDR) < 0.05) responding to changes in CO_2_ at elevated CO_2_ (eCO_2_, 550 ppm) or ambient CO_2_ (aCO_2_, 420 ppm) in *Puccinia sorghi* (*Psor*)-infected plants or mock-inoculated plants at 120 hours post-inoculation (hpi).**

**Supplementary Table 16. Gene ontology (GO) biological process (BP) and molecular function (MF) terms significantly overrepresented in the gene sets developed from clusters of differentially expressed genes (DEG) in response to pathogen infection in plants grown at elevated CO_2_ (eCO_2_, 550 ppm) or ambient CO_2_ (aCO_2_, 420 ppm) conditions and in response to changes in CO_2_ (eCO_2_ vs aCO_2_) in pathogen-infected plants or mock inoculated-plants.** The analysis includes three pathosystems: *Clavibacter nebraskensis* (*Cneb*) at 4 days post-inoculation (dpi), sugarcane mosaic virus (SCMV) at 14 dpi, and *Puccinia sorghi* (*Psor*) at 18, 72, and 120 hours post-inoculation (hpi).

**Supplementary Table 17. Expression in response to pathogen infection in differentially expressed genes (DEG) significant in either (Path eCO_2_ vs Mock eCO_2_) or (Path aCO_2_ vs Mock aCO_2_) and (Mock eCO_2_ vs Mock aCO_2_) or (Path eCO_2_ vs Path aCO_2_).**

**Supplementary Table 18. Expression in response to changes in CO_2_ in differentially expressed genes (DEG) significant in either (Path eCO_2_ vs Mock eCO_2_) or (Path aCO_2_ vs Mock aCO_2_) and (Mock eCO_2_ vs Mock aCO_2_) or (Path eCO_2_ vs Path aCO_2_)**

**Supplementary Table 19. Significantly differentially expressed transcription factors (TF, false discovery rate (FDR) < 0.05) responding to changes in CO_2_ at elevated CO_2_ (eCO_2_, 550 ppm) or ambient CO_2_ (aCO_2_, 420 ppm) in pathogen-infected plants or mock inoculated-plants.** The analysis includes three pathosystems: *Clavibacter nebraskensis* (*Cneb*) at 4 days post-inoculation (dpi), sugarcane mosaic virus (SCMV) at 14 dpi, and *Puccinia sorghi* (*Psor*) at 18, 72, and 120 hours post-inoculation (hpi).

**Supplementary Table 20. Transcription factor binding sites (TFBS) significantly overrepresented (false discovery rate (FDR) < 0.05) among the promoters of differentially expressed genes (DEGs) responding to changes in CO_2_ at elevated CO_2_ (eCO_2_, 550 ppm) or ambient CO_2_ (aCO_2_, 420 ppm) in pathogen-infected plants or mock inoculated-plants.** The analysis includes three pathosystems: *Clavibacter nebraskensis* (*Cneb*) at 4 days post-inoculation (dpi), sugarcane mosaic virus (SCMV) at 14 dpi, and *Puccinia sorghi* (*Psor*) at 18, 72, and 120 hours post-inoculation (hpi).

## References

1. Canadell, J. G. et al. Contributions to accelerating atmospheric CO_2_ growth from economic activity, carbon intensity, and efficiency of natural sinks. Proc. Natl. Acad. Sci. U. S. A. 104, 18866–18870 (2007).

2. Ainsworth, E. A., Sanz-Saez, A. & Leisner, C. P. Crops and rising atmospheric CO_2_: friends or foes? Philos. Trans. R. Soc. Lond. B Biol. Sci. 380, (2025).

3. Meinshausen, M. et al. The shared socio-economic pathway (SSP) greenhouse gas concentrations and their extensions to 2500. Geosci. Model Dev. 13, 3571–3605 (2020).

4. Busch, F. A., Sage, T. L., Cousins, A. B. & Sage, R. F. C3 plants enhance rates of photosynthesis by reassimilating photorespired and respired CO_2_: Intracellular CO_2_ reassimilation. Plant Cell Environ. 36, 200–212 (2013).

5. Leakey, A. D. B. et al. Water use efficiency as a constraint and target for improving the resilience and productivity of C3 and C4 crops. Annu. Rev. Plant Biol. 70, 781–808 (2019).

6. Ainsworth, E. A. & Long, S. P. 30 years of free-air carbon dioxide enrichment (FACE): What have we learned about future crop productivity and its potential for adaptation? Glob. Chang. Biol. 27, 27–49 (2021).

7. Poorter, H. et al. A meta-analysis of responses of C3 plants to atmospheric CO_2_: dose-response curves for 85 traits ranging from the molecular to the whole-plant level. New Phytol. 233, 1560–1596 (2022).

8. Gojon, A., Cassan, O., Bach, L., Lejay, L. & Martin, A. The decline of plant mineral nutrition under rising CO_2_: physiological and molecular aspects of a bad deal. Trends Plant Sci. 28, 185–198 (2023).

9. Hill, A. J. & Shlisel, M. Impact of elevated atmospheric and intercellular CO_2_ on plant defense mechanisms. J. of Crop Health 76, 1307–1315 (2024).

10. Bazinet, Q., Tang, L. & Bede, J. C. Impact of future elevated carbon dioxide on C3 plant resistance to biotic stresses. Mol. Plant. Microbe. Interact. 35, 527–539 (2022).

11. Kazan, K. Plant-biotic interactions under elevated CO_2_: A molecular perspective. Environ. Exp. Bot. 153, 249–261 (2018).

12. Singh, B. K. et al. Climate change impacts on plant pathogens, food security and paths forward. Nat. Rev. Microbiol. 21, 640–656 (2023).

13. Israel, W. K., Watson-Lazowski, A., Chen, Z.-H. & Ghannoum, O. High intrinsic water use efficiency is underpinned by high stomatal aperture and guard cell potassium flux in C3 and C4 grasses grown at glacial CO_2_ and low light. J. Exp. Bot. 73, 1546–1565 (2022).

14. Grulke, N. E. The nexus of host and pathogen phenology: understanding the disease triangle with climate change. New Phytol. 189, 8–11 (2011).

15. Garrett, K. A., Dendy, S. P., Frank, E. E., Rouse, M. N. & Travers, S. E. Climate change effects on plant disease: genomes to ecosystems. Annu. Rev. Phytopathol. 44, 489–509 (2006).

16. Ainsworth, E. A. & Rogers, A. The response of photosynthesis and stomatal conductance to rising [CO_2_]: mechanisms and environmental interactions. Plant Cell Environ. 30, 258–270 (2007).

17. Bredow, M. et al. Elevated CO_2_ alters soybean physiology and defense responses, and has disparate effects on susceptibility to diverse microbial pathogens. New Phytol. 246, 2718–2737 (2025).

18. Foyer, C. H. & Noctor, G. Redox homeostasis and signaling in a higher-CO_2_ world. Annu. Rev. of Plant Bio. 71, 157–182 (2020).

19. Tõldsepp, K. et al. Mitogen-activated protein kinases MPK4 and MPK12 are key components mediating CO_2_-induced stomatal movements. Plant J. 96, 1018–1035 (2018).

20. Li, Z. & Ahammed, G. J. Salicylic acid and jasmonic acid in elevated CO_2_-induced plant defense response to pathogens. J. Plant Physiol. 286, 154019 (2023).

21. Luo, X. et al. Mapping the global distribution of C4 vegetation using observations and optimality theory. Nat. Commun. 15, 1219 (2024).

22. Ranum, P., Peña-Rosas, J. P. & Garcia-Casal, M. N. Global maize production, utilization, and consumption: Maize production, utilization, and consumption. Ann. N. Y. Acad. Sci. 1312, 105–112 (2014).

23. Vaughan, M. M. et al. Interactive effects of elevated [CO_2_] and drought on the maize phytochemical defense response against mycotoxigenic *Fusarium verticillioides*. PLoS One 11, e0159270 (2016).

24. Vaughan, M. M. et al. Effects of elevated [CO_2_] on maize defence against mycotoxigenic *Fusarium verticillioides*. Plant Cell Environ. 37, 2691–2706 (2014).

25. Munkvold, G. P. & White, D. G. PART I: Infectious Diseases. in Compendium of Corn Diseases, Fourth Edition 7–130 (The American Phytopathological Society, 2016).

26. Leakey, A. D. B. et al. Photosynthesis, productivity, and yield of maize are not affected by open-air elevation of CO_2_ concentration in the absence of drought. Plant Physiol. 140, 779–790 (2006).

27. Ruiz-Vera, U. M., Siebers, M. H., Drag, D. W., Ort, D. R. & Bernacchi, C. J. Canopy warming caused photosynthetic acclimation and reduced seed yield in maize grown at ambient and elevated [CO_2_]. Glob. Chang. Biol. 21, 4237–4249 (2015).

28. Osdaghi, E., et al. *Clavibacter nebraskensis* causing Goss’s wilt of maize: Five decades of detaining the enemy in the New World. Mol. Plant Pathol. 24, 675–692 (2023).

29. DeFalco, T. A. & Zipfel, C. Molecular mechanisms of early plant pattern-triggered immune signaling. Mol. Cell 81, 3449–3467 (2021).

30. Kong, L. et al. Dual phosphorylation of DGK5-mediated PA burst regulates ROS in plant immunity. Cell 187, 609–623.e21 (2024).

31. Braidwood, L., Müller, S. Y. & Baulcombe, D. Extensive recombination challenges the utility of Sugarcane mosaic virus phylogeny and strain typing. Sci. Rep. 9, 20067 (2019).

32. Sserumaga, J. P. et al. Identification and diversity of tropical maize inbred lines with resistance to common rust (Schwein). Crop Sci 60, 2971–2989 (2020).

33. Kim, S.-B. et al. Use of the *Puccinia sorghi* haustorial transcriptome to identify and characterize AvrRp1-D recognized by the maize Rp1-D resistance protein. PLoS Pathog 20, e1012662 (2024).

34. Human, M. P., Berger, D. K. & Crampton, B. G. Time-course RNAseq reveals *Exserohilum turcicum* effectors and pathogenicity determinants. Front. Microbiol. 11, 360 (2020).

35. Münch, S. et al. The hemibiotrophic lifestyle of *Colletotrichum* species. J. Plant Physiol. 165, 41–51 (2008).

36. Bickel, J. T. & Koehler, A. M. Review of *Pythium* species causing damping-off in corn. Plant Health Prog. 22, 219–225 (2021).

37. Tian, F., Yang, D.-C., Meng, Y.-Q., Jin, J. & Gao, G. PlantRegMap: charting functional regulatory maps in plants. Nucleic Acids Res. 48, D1104–D1113 (2020).

38. Szklarczyk, D. et al. The STRING database in 2023: protein–protein association networks and functional enrichment analyses for any sequenced genome of interest. Nucleic Acids Res. 51, D638–D646 (2023).

39. Hossain, M. M. et al. Plant disease dynamics in a changing climate: impacts, molecular mechanisms, and climate-informed strategies for sustainable management. Discov. Agric. 2, 132 (2024).

40. Zhou, Y., Vroegop-Vos, I., Schuurink, R. C., Pieterse, C. M. J. & Van Wees, S. C. M. Atmospheric CO_2_ alters resistance of Arabidopsis to *Pseudomonas syringae* by affecting abscisic acid accumulation and stomatal responsiveness to coronatine. Front. Plant Sci. 8, 248465 (2017).

41. Zhou, Y., Van Leeuwen, S. K., Pieterse, C. M. J., Bakker, P. A. H. & Van Wees, S. C. M. Effect of atmospheric CO_2_ on plant defense against leaf and root pathogens of Arabidopsis. Europ. J. Plant Pathol. 154, 31–42 (2019).

42. Hu, Z. et al. High CO_2_- and pathogen-driven expression of the carbonic anhydrase βCA3 confers basal immunity in tomato. New Phytol. 229, 2827–2843 (2021).

43. Li, X. et al. Tomato*-Pseudomonas syringae* interactions under elevated CO_2_ concentration: the role of stomata. J. Exp. Bot. 66, 307–316 (2015).

44. Zhang, S. et al. Antagonism between phytohormone signalling underlies the variation in disease susceptibility of tomato plants under elevated CO_2_. J. Exp. Bot. 66, 1951–1963 (2015).

45. Mhamdi, A. & Noctor, G. High CO_2_ primes plant biotic stress defences through redox-linked pathways. Plant Physiol. 172, 929–942 (2016).

46. Couto, D. & Zipfel, C. Regulation of pattern recognition receptor signalling in plants. Nat. Rev. Immunol. 16, 537–552 (2016).

47. Takahashi, Y. et al. Stomatal CO_2_/bicarbonate sensor consists of two interacting protein kinases, Raf-like HT1 and non-kinase-activity requiring MPK12/MPK4. Sci. Adv. 8, eabq6161 (2022).

48. Vandegeer, R. K., Powell, K. S. & Tausz, M. Barley yellow dwarf virus infection and elevated CO_2_ alter the antioxidants ascorbate and glutathione in wheat. J. Plant Physiol. 199, 96–99 (2016).

49. Trębicki, P. et al. Virus incidence in wheat increases under elevated CO_2_: A 4-year study of yellow dwarf viruses from a free air carbon dioxide facility. Virus Res. 241, 137–144 (2017).

50. Guo, H., Huang, L., Sun, Y., Guo, H. & Ge, F. The contrasting effects of elevated CO_2_ on TYLCV infection of tomato genotypes with and without the resistance gene, Mi-1.2. Front. Plant Sci. 7, 1680 (2016).

51. Matros, A. et al. Growth at elevated CO_2_ concentrations leads to modified profiles of secondary metabolites in tobacco cv. SamsunNN and to increased resistance against infection with potato virus Y. Plant Cell Environ. 29, 126–137 (2006).

52. Ye, L., Fu, X. & Ge, F. Elevated CO_2_ alleviates damage from Potato virus Y infection in tobacco plants. Plant Sci. 179, 219–224 (2010).

53. Scandolera, T. et al. Effects of elevated CO_2_ on bean pod mottle virus infection in both incompatible and compatible interactions with *Phaseolus vulgaris* L. Plant Cell Environ. 0, 1–20 (2025).

54. Carbonell, A. & Carrington, J. C. Antiviral roles of plant ARGONAUTES. Curr. Opin. Plant Biol. 27, 111–117 (2015).

55. Fang, X. & Qi, Y. RNAi in plants: An argonaute-centered view. Plant Cell 28, 272–285 (2016).

56. Mahindapala, R. Host and environmental effects on the infection of maize by *Puccinia sorghi*. Ann. Appl. Biol. 89, 411–416 (1978).

57. Mapuranga, J., Zhang, N., Zhang, L., Chang, J. & Yang, W. Infection strategies and pathogenicity of biotrophic plant fungal pathogens. Front. Microbiol. 13, 799396 (2022).

58. Helfer, S. Rust fungi and global change. New Phytol. 201, 770–780 (2014).

59. Porras, R., Miguel-Rojas, C., Lorite, I. J., Pérez-de-Luque, A. & Sillero, J. C. Characterization of durum wheat resistance against leaf rust under climate change conditions of increasing temperature and [CO_2_]. Sci. Rep. 13, 22001 (2023).

60. Xu, Z., Jiang, Y., Jia, B. & Zhou, G. Elevated-CO_2_ response of stomata and its dependence on environmental factors. Front. Plant Sci. 7, 657 (2016).

61. Mikkelsen, B. L., Olsen, C. E. & Lyngkjær, M. F. Accumulation of secondary metabolites in healthy and diseased barley, grown under future climate levels of CO_2_, ozone and temperature. Phytochemistry 118, 162–173 (2015).

62. Kobayashi, T. et al. Effects of elevated atmospheric CO_2_ concentration on the infection of rice blast and sheath blight. Phytopathology 96, 425–431 (2006).

63. Frew, A. & Price, J. N. Mycorrhizal-mediated plant–herbivore interactions in a high CO_2_ world. Funct. Ecol. 33, 1376–1385 (2019).

64. Jwa, N.-S. & Walling, L. L. Influence of elevated CO_2_ concentration on disease development in tomato. New Phytol. 149, 509–518 (2001).

65. Bao, Y. et al. Assessing plant performance in the Enviratron. Plant Methods 15, 117 (2019).

66. Jaggard, K. W., Qi, A. & Ober, E. S. Possible changes to arable crop yields by 2050. Philos. Trans. R. Soc. Lond. B Biol. Sci. 365, 2835–2851 (2010).

67. Ceulemans, R., VAN Praet, L. & Jiang, X. N. Effects of CO_2_ enrichment, leaf position and clone on stomatal index and epidermal cell density in poplar (*Populus*). New Phytol. 131, 99–107 (1995).

68. Sultana, S. N., et al. Optimizing the experimental method for stomata-profiling automation of soybean leaves based on deep learning. Plants 10, (2021).

69. Eisele, J. F., Fäßler, F., Bürgel, P. F. & Chaban, C. A rapid and simple method for microscopy-based stomata analyses. PLoS One 11, e0164576 (2016).

70. Shu, Y., Wan-Ting, J., Ya-Ning, Y. & Yu-Han, F. An optimized CTAB method for genomic DNA extraction from freshly-picked pinnae of fern, L. Bio Protoc 8, e2906 (2018).

71. de Sena Brandine, G. & Smith, A. D. Falco: high-speed FastQC emulation for quality control of sequencing data. F1000Res. 8, 1874 (2019).

72. Dobin, A. et al. STAR: ultrafast universal RNA-seq aligner. Bioinformatics 29, 15–21 (2013).

73. Li, H. et al. The sequence alignment/map format and SAM tools. Bioinformatics 25, 2078–2079 (2009).

74. RStudio: Integrated Development Environment for R. (Posit Software, Boston, MA, 2024).

75. Lawrence, M. et al. Software for computing and annotating genomic ranges. PLoS Comput. Biol. 9, e1003118 (2013).

76. Wickham, H. Ggplot2: elegant graphics for data analysis. (Springer, New York, NY, 2009).

77. Robinson, M. D., McCarthy, D. J. & Smyth, G. K. edgeR: a Bioconductor package for differential expression analysis of digital gene expression data. Bioinformatics 26, 139–140 (2010).

78. Murtagh, F. Multidimensional Clustering Algorithms. (Springer, New York, NY, 1985).

79. Camacho, C. et al. BLAST+: architecture and applications. BMC Bioinformatics 10, 421 (2009).

80. Suzek, B. E., Huang, H., McGarvey, P., Mazumder, R. & Wu, C. H. UniRef: comprehensive and non-redundant UniProt reference clusters. Bioinformatics 23, 1282–1288 (2007).

81. Ashburner, M. et al. Gene ontology: tool for the unification of biology. The Gene Ontology Consortium. Nat. Genet. 25, 25–29 (2000).

82. Bonferroni, C. E. Il calcolo delle assicurazioni su gruppi di teste. Studi onore del professore salvatore ortu carboni. 13–60 (1935).

83. Bredow, M., Sementchoukova, I., Siegel, K. & Monaghan, J. Pattern-triggered oxidative burst and seedling growth inhibition assays in *Arabidopsis thaliana*. J. Vis. Exp. 147, e59437 (2019).

84. Wang, J. et al. A regulatory module controlling homeostasis of a plant immune kinase. Mol. Cell 69, 493–504.e6 (2018).

85. Balcke, G. U. et al. An UPLC-MS/MS method for highly sensitive high-throughput analysis of phytohormones in plant tissues. Plant Methods 8, 47 (2012).

86. Bligh, E. G. & Dyer, W. J. A rapid method of total lipid extraction and purification. Can. J. Biochem. Physiol. 37, 911–917 (1959).

87. Triebl, A. et al. Quantitation of phosphatidic acid and lysophosphatidic acid molecular species using hydrophilic interaction liquid chromatography coupled to electrospray ionization high resolution mass spectrometry. J. Chromatogr. A 1347, 104–110 (2014).

88. Smith, C. A. et al. METLIN: a metabolite mass spectral database. Ther. Drug Monit. 27, 747–751 (2005).

89. Vidaver, A. K. Synthetic and complex media for the rapid detection of fluorescence of phytopathogenic pseudomonads: effect of the carbon source. Appl. Microbiol. 15, 1523–1524 (1967).

90. Hu, Y. et al. Analysis of extreme phenotype bulk copy number variation (XP-CNV) identified the association of rp1 with resistance to goss’s wilt of maize. Front. Plant Sci. 9, 110 (2018).

91. Mbofung, G. C. Y., Sernett, J., Horner, H. T. & Robertson, A. E. Comparison of susceptible and resistant maize hybrids to colonization by *Clavibacter michiganensis* subsp. *nebraskensis*. Plant Dis. 100, 711–717 (2016).

92. Hwang, I. S., Oh, E.-J., Kim, D. & Oh, C.-S. Multiple plasmid-borne virulence genes of *Clavibacter michiganensis* ssp. *capsici* critical for disease development in pepper. New Phytol. 217, 1177–1189 (2018).

93. Simko, I. & Piepho, H.-P. The area under the disease progress stairs: calculation, advantage, and application. Phytopathology 102, 381–389 (2012).

94. Skoppek, C. I. & Streubel, J. Simplifying barley leaf rust research: an easy and reproducible infection protocol for on a small laboratory scale. Bio Protoc 13, e4721 (2023).

95. Reuveni, R., Agapov, V. & Reuveni, M. Foliar spray of phosphates induces growth increase and systemic resistance to *Puccinia sorghi* in maize. Plant Pathol. 43, 245–250 (1994).

96. Holan, K. L., White, C. H. & Whitham, S. A. Application of a U-Net neural network to the *Puccinia sorghi*-maize pathosystem. Phytopathology 114, 990–999 (2024).

97. Sermons, S. M. & Balint-Kurti, P. J. Large scale field inoculation and scoring of maize southern LeafBlight and other maize foliar fungal diseases. Bio Protoc. 8, e2745 (2018).

98. Vieira, R. A. et al. A new diagrammatic scale for the assessment of northern corn leaf blight. Crop Prot. 56, 55–57 (2014).

99. Belisário, R., Robertson, A. E. & Vaillancourt, L. J. Maize anthracnose stalk rot in the genomic era. Plant Dis. 106, 2281–2298 (2022).

100. Christensen, J. J., Wilcoxson, R. D., American Phytopathological Society & Christensen, J. J. Stalk Rot of Corn. (American Phytopathological Society, 1966).

101. Zhang, B. Q. & Yang, X. B. Pathogenicity of *Pythium* populations from corn-soybean rotation fields. Plant Dis. 84, 94–99 (2000).

102. Broders, K. D., Lipps, P. E., Paul, P. A. & Dorrance, A. E. Characterization of *Pythium* spp. associated with corn and soybean seed and seedling disease in Ohio. Plant Dis. 91, 727–735 (2007).

103. Langenhoven, B., Murray, S. L. & Crampton, B. G. Quantitative detection of *Exserohilum turcicum* in northern leaf blight diseased sorghum and maize leaves. Australas. Plant Pathol. 49, 609–617 (2020).

104. Weihmann, F. et al. Correspondence between symptom development of *Colletotrichum graminicola* and fungal biomass, quantified by a newly developed qPCR assay, depends on the maize variety. BMC Microbiol. 16, 94 (2016).

105. Cooke, D. E., Drenth, A., Duncan, J. M., Wagels, G. & Brasier, C. M. A molecular phylogeny of *Phytophthora* and related oomycetes. Fungal Genet. Biol. 30, 17–32 (2000).

106. Lin, Y. et al. Validation of potential reference genes for qPCR in maize across abiotic stresses, hormone treatments, and tissue types. PLoS One 9, e95445 (2014).

107. Yan, Y. et al. Disruption of *OPR7* and *OPR8* reveals the versatile functions of jasmonic acid in maize development and defense. Plant Cell 24, 1420–1436 (2012).

