## Supplementary Figures for "Effects of atmospheric CO_2_ levels on the susceptibility of maize to diverse pathogens"

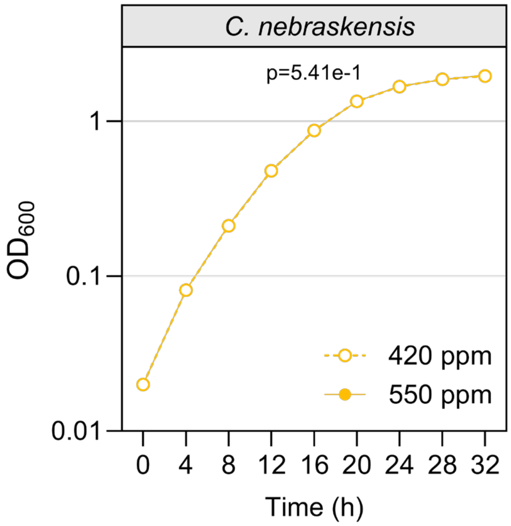
C. nebraskensis
p=5.41e-1
1
0.1
0.01
OD600
420 ppm
550 ppm
0
4
8
12
16
20
24
28
32
Time (h)

**a**

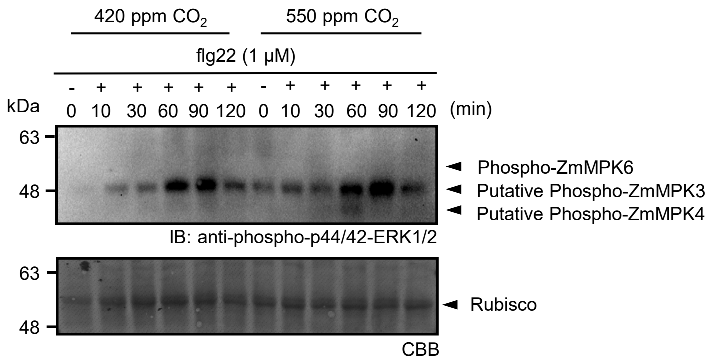
420 ppm $CO_2$
550 ppm $CO_2$
flg22 (1 μM)
- + + + + + - + + + + +
kDa
0 10 30 60 90 120 0 10 30 60 90 120 (min)
63
48
Phospho-ZmMPK6
Putative Phospho-ZmMPK3
Putative Phospho-ZmMPK4
IB: anti-phospho-p44/42-ERK1/2
63
48
Rubisco
CBB

**c**

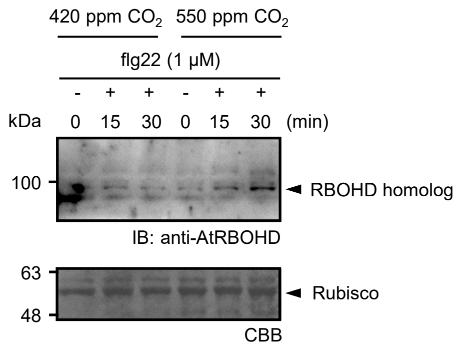
420 ppm $CO_2$
550 ppm $CO_2$
flg22 (1 μM)
- + + - + +
kDa
0 15 30 0 15 30 (min)
100
RBOHD homolog
IB: anti-AtRBOHD
63
48
Rubisco
CBB

**b**

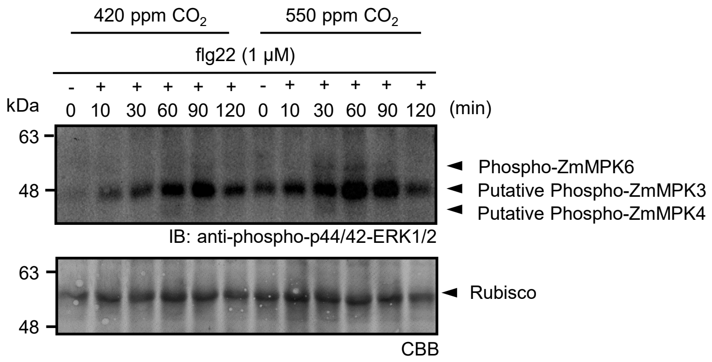
420 ppm $CO_2$
550 ppm $CO_2$
flg22 (1 μM)
- + + + + + - + + + + +
kDa
0 10 30 60 90 120 0 10 30 60 90 120 (min)
63
48
Phospho-ZmMPK6
Putative Phospho-ZmMPK3
Putative Phospho-ZmMPK4
IB: anti-phospho-p44/42-ERK1/2
63
48
Rubisco
CBB

**d**

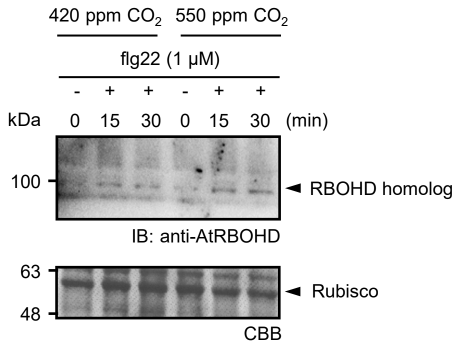
420 ppm $CO_2$
550 ppm $CO_2$
flg22 (1 μM)
- + + - + +
kDa
0 15 30 0 15 30 (min)
100
RBOHD homolog
IB: anti-AtRBOHD
63
48
Rubisco
CBB

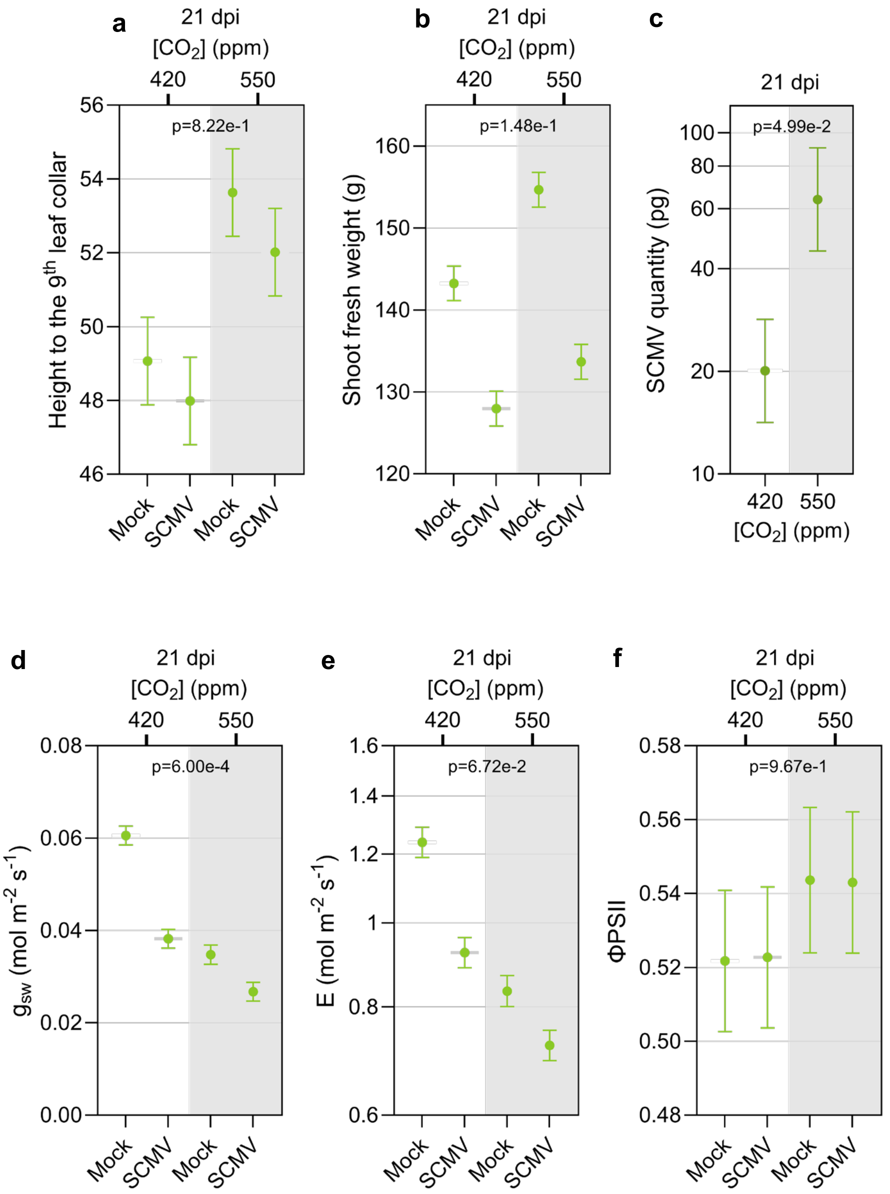
a
21 dpi
[CO2] (ppm)
420
550
p=8.22e-1
Height to the 9th leaf collar
56
54
52
50
48
46
Mock
SCMV
Mock
SCMV
b
21 dpi
[CO2] (ppm)
420
550
p=1.48e-1
Shoot fresh weight (g)
160
150
140
130
120
Mock
SCMV
Mock
SCMV
c
21 dpi
p=4.99e-2
SCMV quantity (pg)
100
80
60
40
20
10
420
550
[CO2] (ppm)
d
21 dpi
[CO2] (ppm)
420
550
p=6.00e-4
gsw (mol m-2 s-1)
0.08
0.06
0.04
0.02
0.00
Mock
SCMV
Mock
SCMV
e
21 dpi
[CO2] (ppm)
420
550
p=6.72e-2
E (mol m-2 s-1)
1.6
1.4
1.2
1
0.8
0.6
Mock
SCMV
Mock
SCMV
f
21 dpi
[CO2] (ppm)
420
550
p=9.67e-1
ΦPSII
0.58
0.56
0.54
0.52
0.50
0.48
Mock
SCMV
Mock
SCMV

**Supplementary Fig. 4**

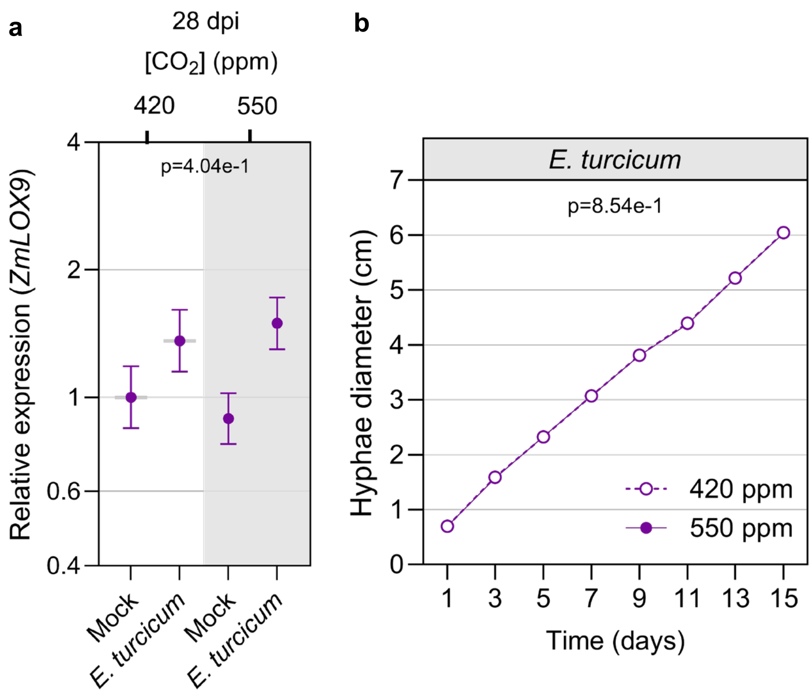

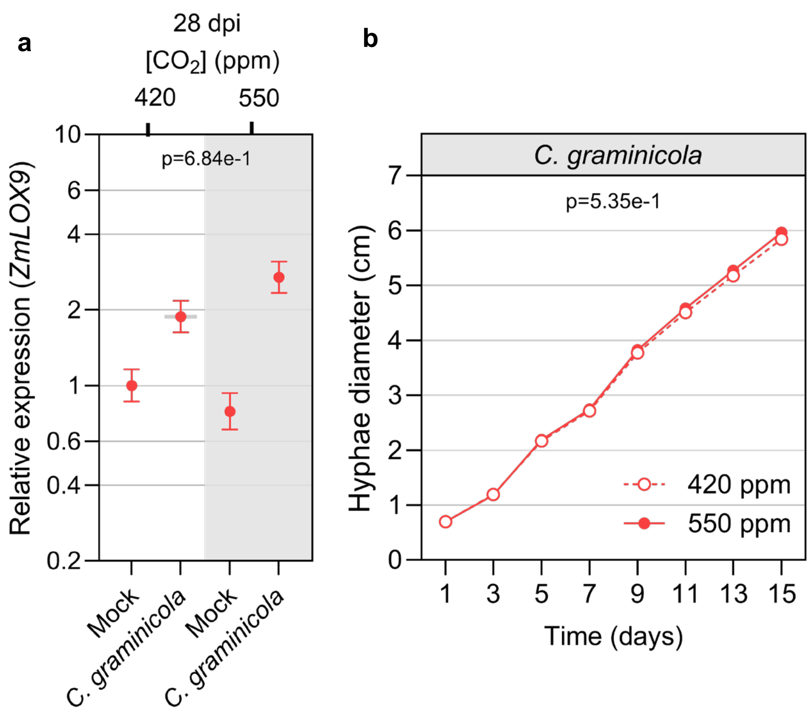

a
28 dpi
[CO2] (ppm)
420
550
p=6.84e-1
Relative expression (ZmLOX9)
10
6
4
2
1
0.6
0.4
0.2
Mock
C. graminicola
Mock
C. graminicola
b
C. graminicola
p=5.35e-1
Hyphae diameter (cm)
7
6
5
4
3
2
1
0
420 ppm
550 ppm
1
3
5
7
9
11
13
15
Time (days)

**Supplementary Fig. 6**

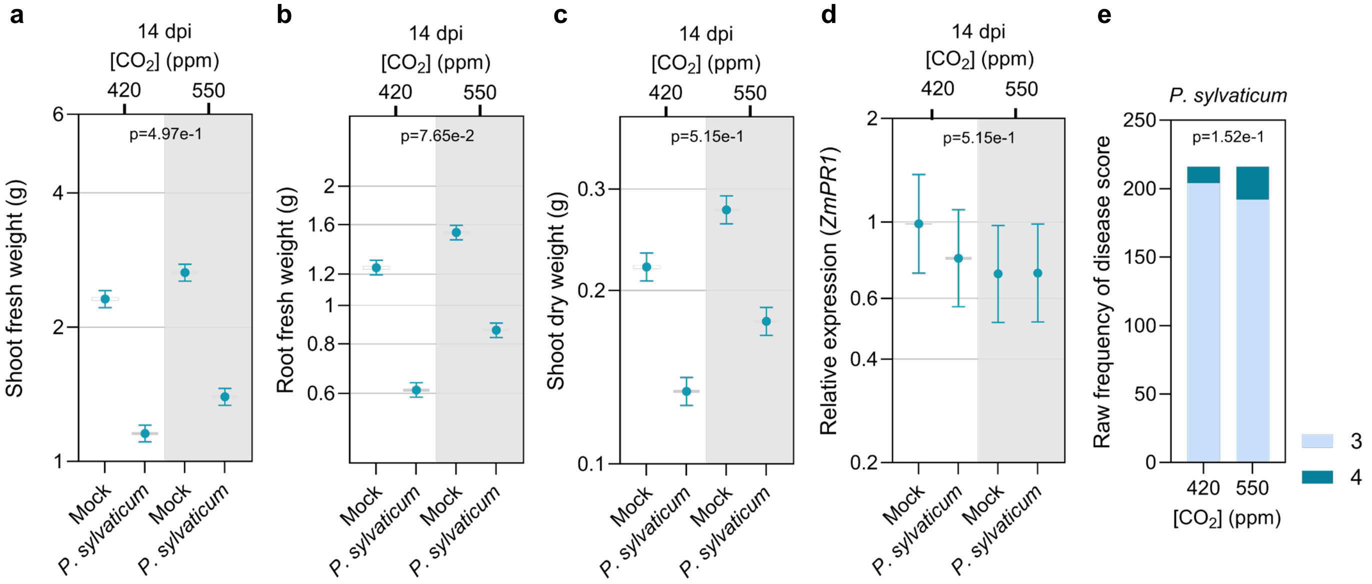

**Supplementary Fig. 7**

**a**

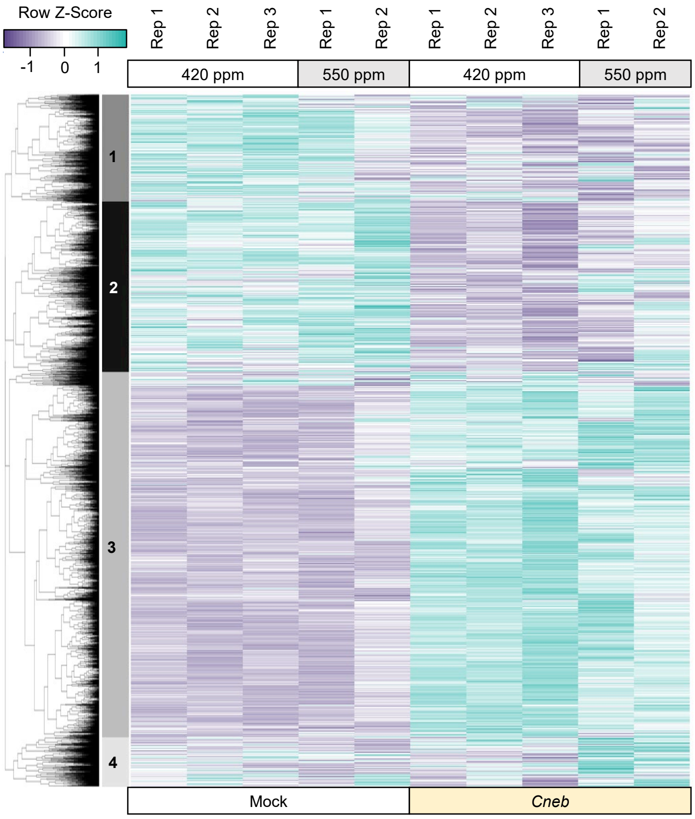


**b**

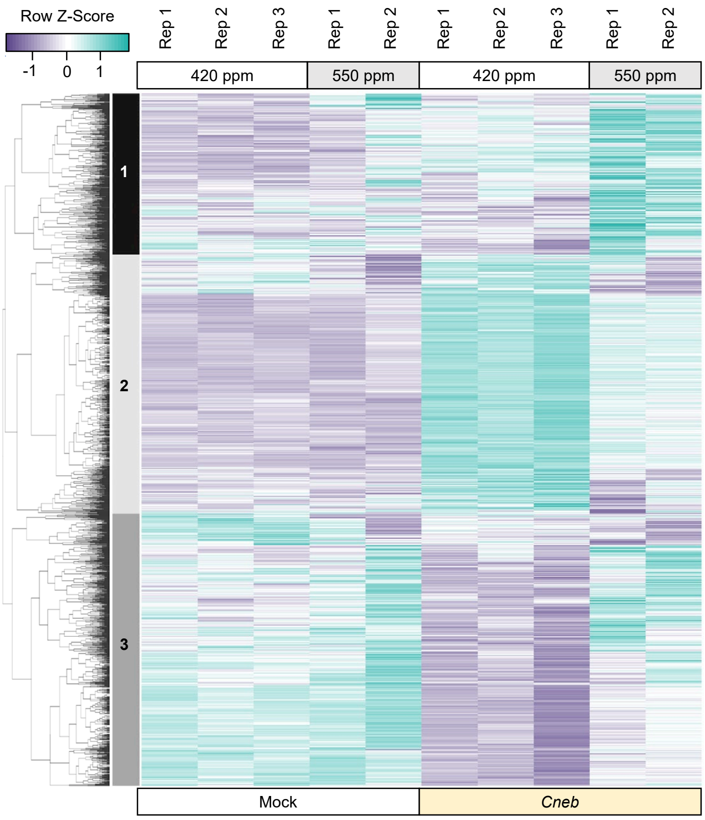

**Supplementary Fig. 8**

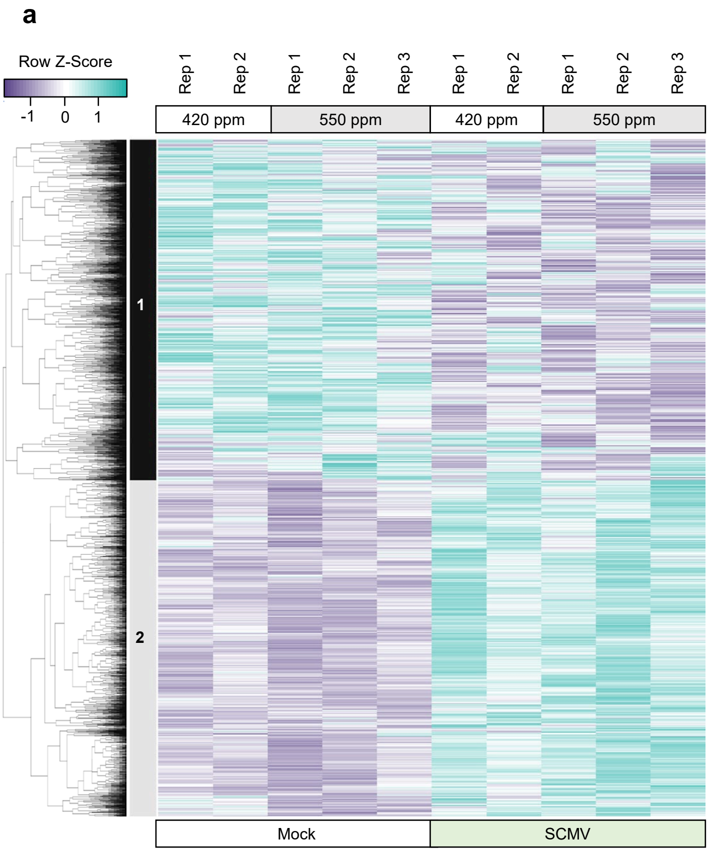

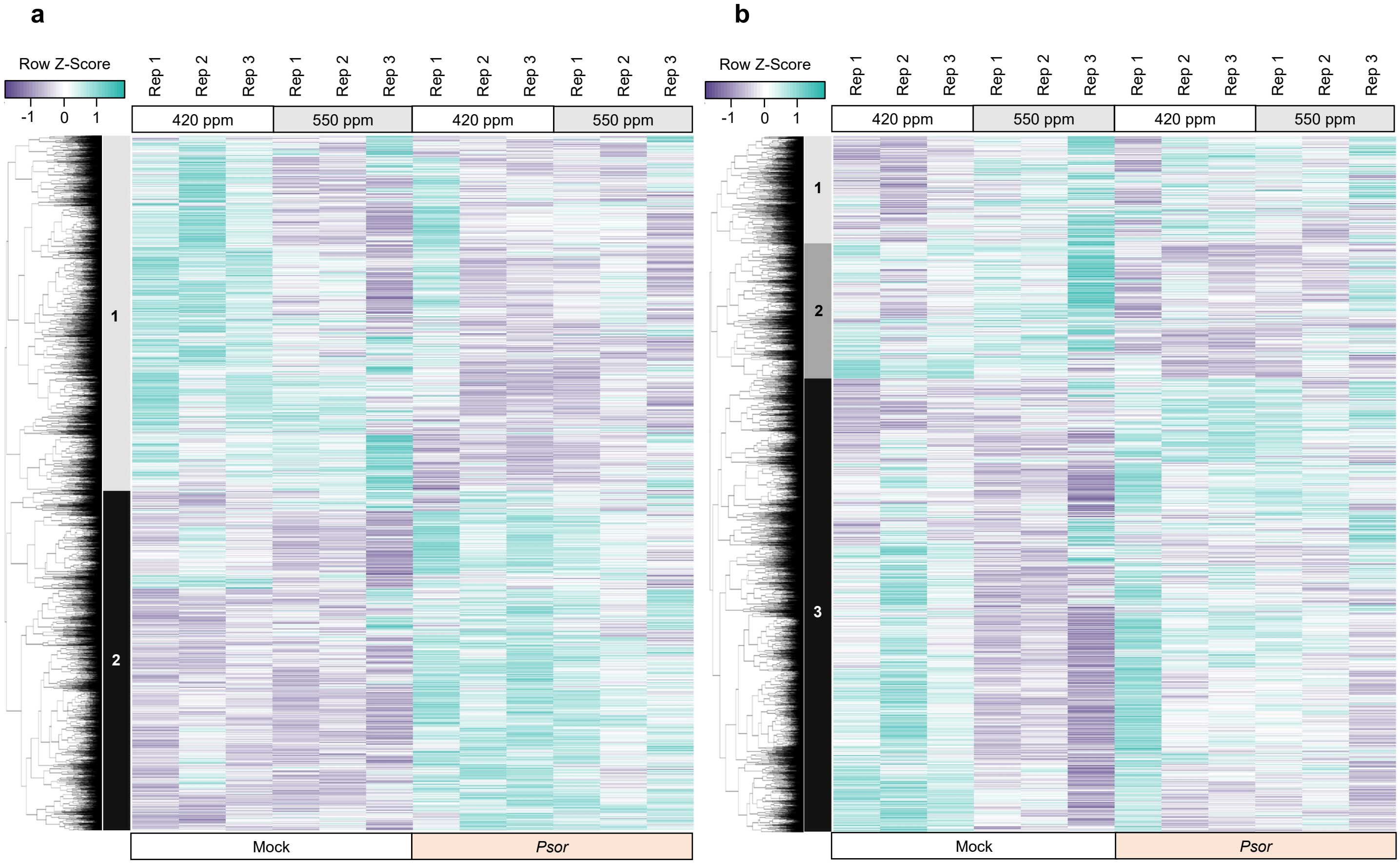
a
Row Z-Score
-1
0
1
Rep 1
Rep 2
Rep 3
420 ppm
550 ppm
1
2
Mock
Psor
b
Row Z-Score
3

**Supplementary Fig. 10**

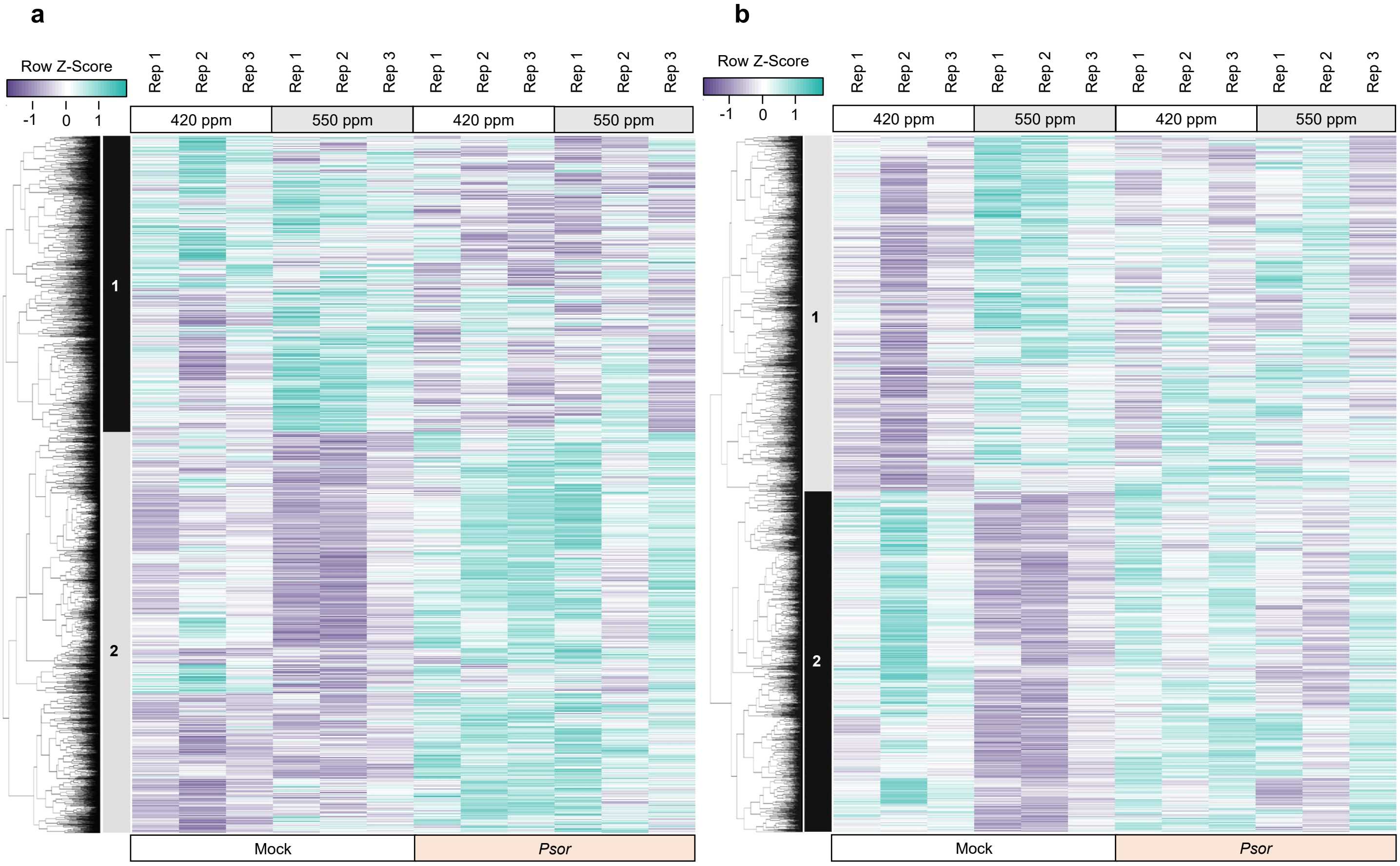

**Supplementary Fig. 11**

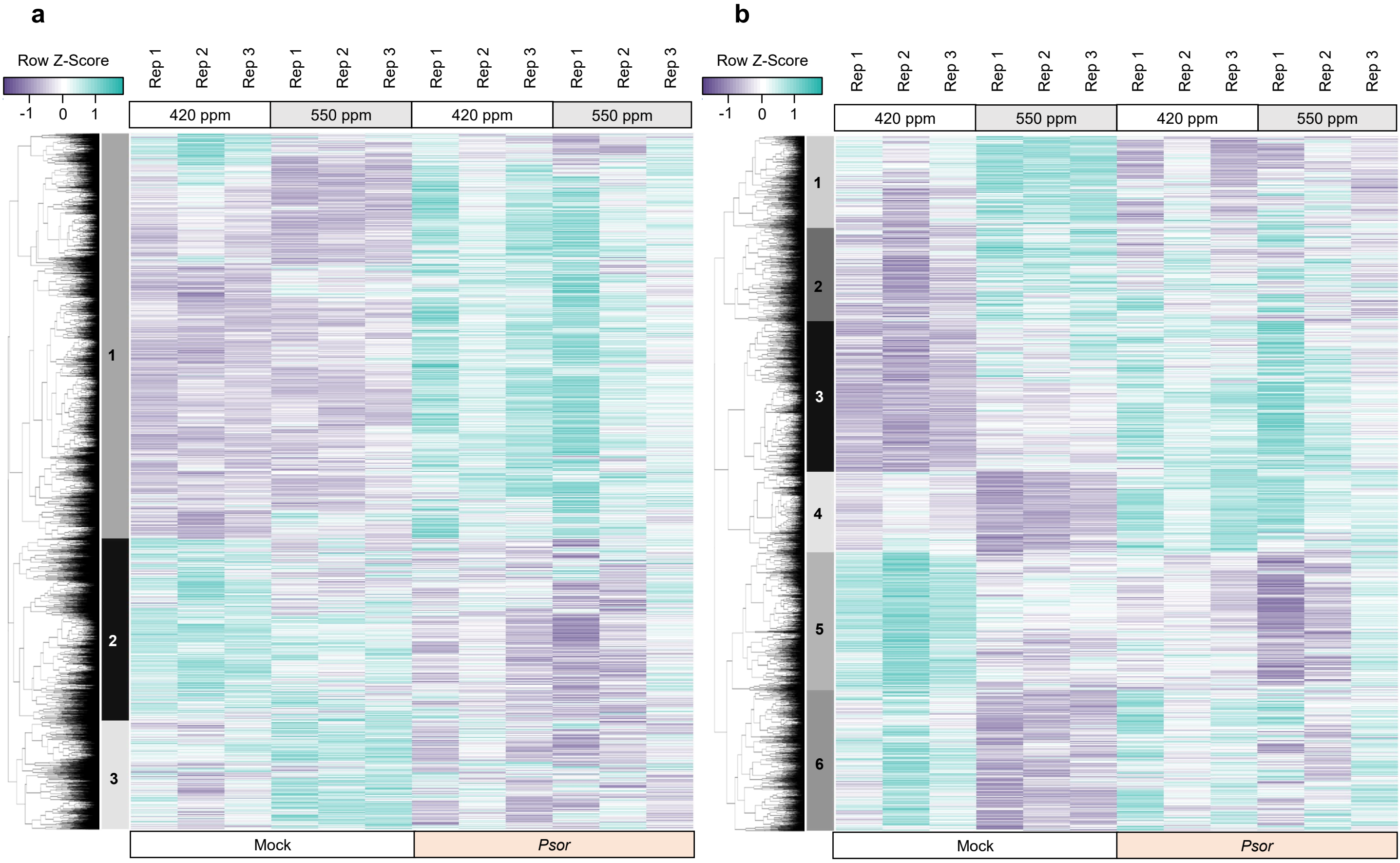
